# Adaptive motility and feeding strategies of deep-branching predatory protists

**DOI:** 10.64898/2026.10.08.757618

**Authors:** Julien Bouvard, Angéline Girard, Samuel A. Bentley, Harold Auradou, Kirsty Y. Wan

## Abstract

Heterotrophic flagellates play a major role in nutrient recycling and shaping aquatic microbial communities. Many motile species have evolved diverse predatory strategies to exploit microbial prey, from bacteria to small eukaryotes. Excavates, deep-branching eukaryotic lineages with distinctive biflagellate morphology and conserved cytoskeletal features, exhibit varied feeding behaviours, including grazing, raptorial feeding, and filter-feeding. However, the temporal dynamics of their motility, feeding, and spatial distribution in relation to bacterial population density remain under-explored. Here, we quantify the behavioural ecology of excavates using high-resolution, single-cell tracking and quantitative analysis. Focussing on two Discoba species – the marine *Jakoba libera* and freshwater *Reclinomonas americana* – we identify different behavioural patterns: *J. libera* swims, forages and feeds, while *R. americana* swims and feeds. Prey availability differentially affects motility: *J. libera* increases the number of swimming cells when food is scarce, while *R. americana* shows the opposite trend. Spatial statistics reveal fundamental differences in their feeding and redistribution strategies. *J. libera* acts as an active forager, whereas *R. americana* behaves as a strict filter feeder. A third species, the anaerobic *Kipferlia bialata* (from a different clade), adopts a foraging strategy similar to *J. libera*. Together, these results elucidate the adaptive relationship between prey availability, feeding modes, and motility in bacteriovorous excavate flagellates, offering new insights into the ecological and evolutionary dynamics of microbial predators.

**SIGNIFICANCE:** Microbial hunters and predators that feed on other small organisms play an important role in aquatic food webs. Many species are motile, with seemingly adaptive behaviours that are difficult to quantify. Here, we focus on different species of early-diverging biflagellates that share a common morphology consisting of two flagella and a feeding groove, which it uses to feed on surface-associated bacterial prey. We demonstrate how the predator’s behavioural profiles shift as prey abundance changes over many days of a culture cycle, accompanied by evolving patterns of spatial distribution. Species-specific differences provide insights into the mechanisms of environmental adaptation in early eukaryotes.

---

Microbial eukaryotes, together with fungi, bacteria, and archaea, form complex communities that display an immense cryptic biodiversity that shapes ecological dynamics, food web interactions, and ecosystem responses to environmental change [1, 2]. Among these, heterotrophic flagellates, single cells that use whip-like flagella to swim and engulf other organisms as food, are key consumers of bacteria and other small prey [3]. They recycle nutrients and channel energy up through the higher trophic levels, and are thus critical for ecosystem structure and functioning across marine and freshwater systems. In the oceans, microbial communities dominate biomass and regulate biogeochemical cycles, contributing significantly to global carbon fixation and nutrient cycling [4–6]. Similarly, freshwater ecosystems harbour vast assemblages of microscopic organisms that underlie key processes in lakes, rivers, and reservoirs [7, 8].

Over time, distinct environmental pressures have driven the evolution of substantial diversity and behavioural sophistication in the feeding, foraging and motility habits of heterotrophic flagellates [9–12]. Motility is essential to locate food, nutrients, or to escape toxic environments and evade predators [13, 14]. Adaptive motility, including responses to chemical [15], light [16, 17], mechanical [18–20], as well as biotic cues [21, 22], is central to their ecological success. Many species are predatory and use their flagella to actively hunt and consume their prey [23]. Some even have specialised feeding or prey-capture apparatuses to efficiently exploit microbial prey in spatially heterogenous environments, particularly on or near surfaces [24].

Among eukaryotic supergroups, free-living excavates and other excavate-like organisms have a remarkable functional diversity [25], including a heterotrophic (indeed phagotrophic) lifestyle. They are hypothesised to be the earliest-diverging lineages and retain many features of the Last Common Eukaryotic Ancestor (LECA) [26, 27]. Some feed on surface-associated bacteria, generating feeding currents while phagocytosing captured prey [10]. Others are instead eukaryovorous and engage in complex prey handling processes to feed on other eukaryotes that may be almost as large as the excavate itself [28]. Yet, despite their ecological importance, little is known about the detailed feeding and motility strategies of excavates. Studying these processes would provide key insights into how complex behavioural traits such as predation evolved and diversified in early unicellular eukaryotes [11],

Here, we explore how excavate predators dynamically adapt their feeding and motility behaviours to changing environmental conditions, going well beyond descriptive accounts. We focus on ‘typical excavates’, a paraphyletic subgroup with species drawn from three phylogenetic groups: Discoba, Metamonada and Malawimonadida [25]. These cells have a distinctive morphology [27], comprising two heterodynamic flagella and a ventral feeding groove, see Fig. 1a. The posterior flagellum beats within the groove, while the anterior flagellum beats far from the cell body. Small bacterial prey are ingested via a cytopharynx located at the posterior end of the ventral groove [24]. We created a framework to systematically monitor and quantify the adaptive foraging strategies of excavates during typical batch culture progression, from resource-rich to resource-limited phases. Our approach is multiscale, with high frame-rate, high resolution imaging to capture single-cell behaviour, and lower frame rate and magnification to perform population scale tracking and automated image analysis.

**Figure 1.**
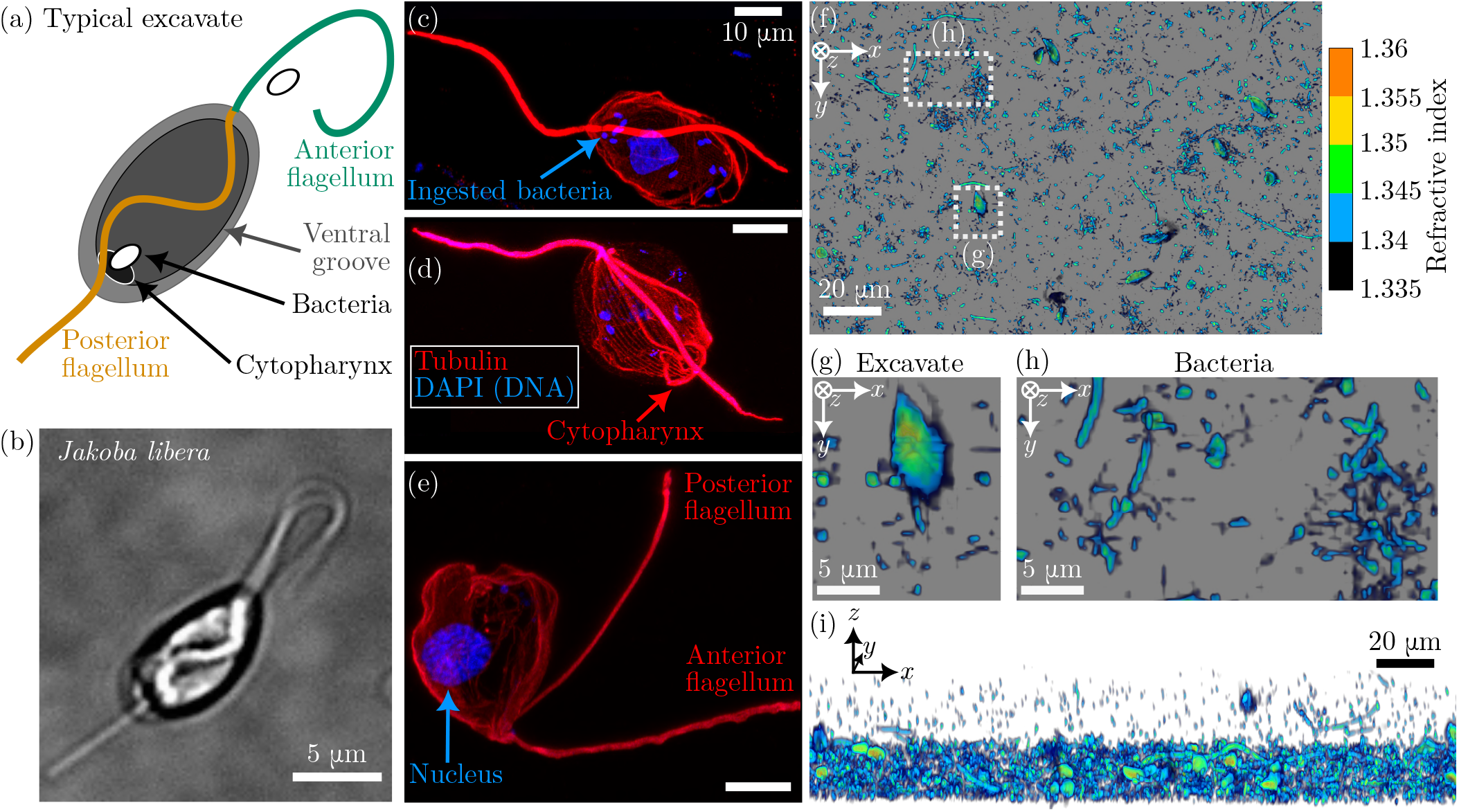
Morphology and habitat of single-celled predatory flagellates known as excavates. (a) Schematic of a typical excavate cell, view from the ventral side. (b) Bright-field image of *Jakoba libera* swimming in the bulk, taken at 101× magnification. (c-e) Maximum intensity projection of Ultrastructure Expansion Microscopy (U-ExM) images: (c) *J. libera*, (d) *Kipferlia bialata*, and (e) *Reclinomonas americana*. Cells were respectively expanded 4.0×, 3.5× and 4.2× before imaging. [Tubulin (red); DAPI (blue)]. Scalebars: 10 µm in all three panels. (f-i) Holotomography image of *K. bialata* excavates in a bacteria-rich environment close to a surface, colour-coded by refractive index: (f) bottom-view, (g-h) zooms of bottom-view, and (i) slightly tilted side view.

Investigating three species from two different clades: the aerobic marine *Jakoba libera* and aerobic fresh-water *Reclinomonas americana* from Discoba [29, 30], and the anaerobic marine species *Kipferlia bialata* from Metamonada [31], we show how an apparently similar morphology can lead to starkly contrasting behaviours. While some species behave as active foragers, others instead only perform filter feeding while attached surfaces. Three distinct modes are identified: feeding on surfaces, swimming in the bulk, and a novel, previously undescribed foraging behaviour along surfaces. Excavates can shift between these behaviours, depending on environmental conditions. We classify these detailed behaviours and quantify how the predator’s behavioural profile shifts across a vast dynamic range of bacteria densities, over many days of a culture cycle. Finally, we develop spatial statistics tools to measure changes in the surface distribution of excavates over time, again highlighting key differences between the forager *J. libera* and the filter feeder *R. americana*. Our data is used to determine the presence of short-range attractive and medium-range repulsive interactions, which could lead to self-organised structures at different scales, including hyperuniform states [32, 33]. Together, our results reveal how bacteriovorous flagellates adapt and transition between different behavioural strategies depending on resource availability in a species-dependent manner, and provide a new foundation for systematic comparative studies of behaviour and sensory processing at the microscopic scale.

## RESULTS

### A. Visualisation of typical excavates & their prey

When imaged close to surfaces, individual excavates appear distinctly with both flagella and ventral feeding groove visible (see Fig. 1b). To visualise cell morphology, we conduct Ultrastructure Expansion Microscopy (U-ExM) imaging (see Methods) [34, 35]. U-ExM expands the cell ≈ 4 × while preserving the relative organisation of internal structures, which enhances the resolution of cellular morphology and allows us to obtain quantitative parameters such as feeding groove dimensions (Table S2). Tubulin (*α* and *β*-tubulin) and DNA (DAPI) were fluorescently labelled in the samples, and maximum intensity projection images of confocal z-stacks were obtained for each species (Fig. 1c-e). The two flagella, whose axonemes are composed of microtubules, are clearly visible, along with other cytoskeletal filaments that define the morphology of the cell body.

Notable differences exist between the three species. In *R. americana*, the cell body appears nebulous and less defined in the tubulin channel, whereas *K. bialata* and *J. libera* exhibit distinct arrays of microtubules beneath the membrane cortex. In *K. bialata*, the cytopharynx (i.e., the “mouth”) manifests as a ring of microtubules located at the posterior end of the cell, see Fig. 1d. Morphological measurements from the U-ExM images are consistent with literature values obtained from other imaging modalities [10, 36]. The DNA staining enables visualisation of the nuclei of both predator and ingested bacterial prey. The nucleus of the eukaryotic cell is typically observed as a large, well-defined structure. This nucleus can appear either alone (as in Fig. 1e) or alongside multiple DNA spots, which represent ingested bacteria (Fig. 1c-d). For *K. bialata*, ingested bacteria are observed at the posterior end of the cell near the cytopharynx, while for *J. libera* they also appear at the anterior end (additional examples are provided in Fig. S1).

Excavates occupy environments rich in surfaces [10], including marine [29, 31] and freshwater sediments [30]. They were also only observed to feed on or near surfaces, which prompted us to examine the microbial composition of surfaces present in our cultures. Here the microenvironment is very diverse, with bacteria of different sizes, shapes, and motility (see Fig. S2 for snapshots). Both sessile and motile bacteria are abundant in *K. bialata* cultures, whereas most of the bacteria found in *J. libera* and *R. americana* cultures are sessile.

To visualise the natural habitat structure of typical excavates under the conditions they are cultured in the laboratory, we imaged the bottom surface of one-week-old samples by holotomography (see Methods). Holotomography is a label-free imaging technique that uses refractive index variations within a specimen, allowing for a non-invasive 3D reconstruction of the surface topography and species composition of the suspension. As the refractive indices of bacteria and excavates are both slightly higher than that of water (1.3337), they are clearly resolved by holotomography. Refractive indices of the bacteria varied from 1.335 to 1.35, for the excavates between 1.34 and 1.355. Fig. 1f-i shows one example of 2D projections of 3D holotomography from a *K. bialata* sample. With the selected color map, bacteria appear as blue dots with tints of black and green, whereas the excavates appear green with shades of yellow and blue. We deduce that the excavates encounter a stratified environment consisting of a heterogeneous biofilm containing a very high density of bacteria on the lower surface, especially for *K. bialata*, with a decreasing density of bacteria extending into the water column.

### B. Diversity of motility behaviours

We turn to the distinct motility and behavioural repertoires of typical excavates, focussing on *J. libera* and *R. americana*. The third species, *K. bialata*, shows qualitatively similar behaviours as *J. libera* (see Fig. S5). While both *J. libera* and *R. americana* belong to the Discoba group, they inhabit different natural environments as the former is marine and the latter is freshwater.

Detailed observations of motility trajectories allow us to classify the behaviour of excavates into discrete modes, as done previously in many other microorganisms [37–39]. Three distinct behaviours were found in *J. libera*: swimming in the bulk, foraging along surfaces, and feeding on surfaces, see Fig. 2a. Dense *J. libera* suspensions were recorded with a 10× objective (see Methods), with images taken close to the bottom surface to capture all behaviours simultaneously, with d*t* = 1 s time-steps. For simplicity, each trajectory was classified as one of the three behaviours: feeding, foraging, or swimming (see Methods). The mean squared displacement (MSD) was calculated for each sub-population, see Fig. 2b. The MSD of the swimming population (green circles) follows a *t*^2^ trend, characteristic of ballistic behaviour, while the MSD of the foraging population is proportional to *t*, characteristic of diffusive behaviour. A sub-diffusive behaviour is observed for the feeding sub-population.

**Figure 2.**
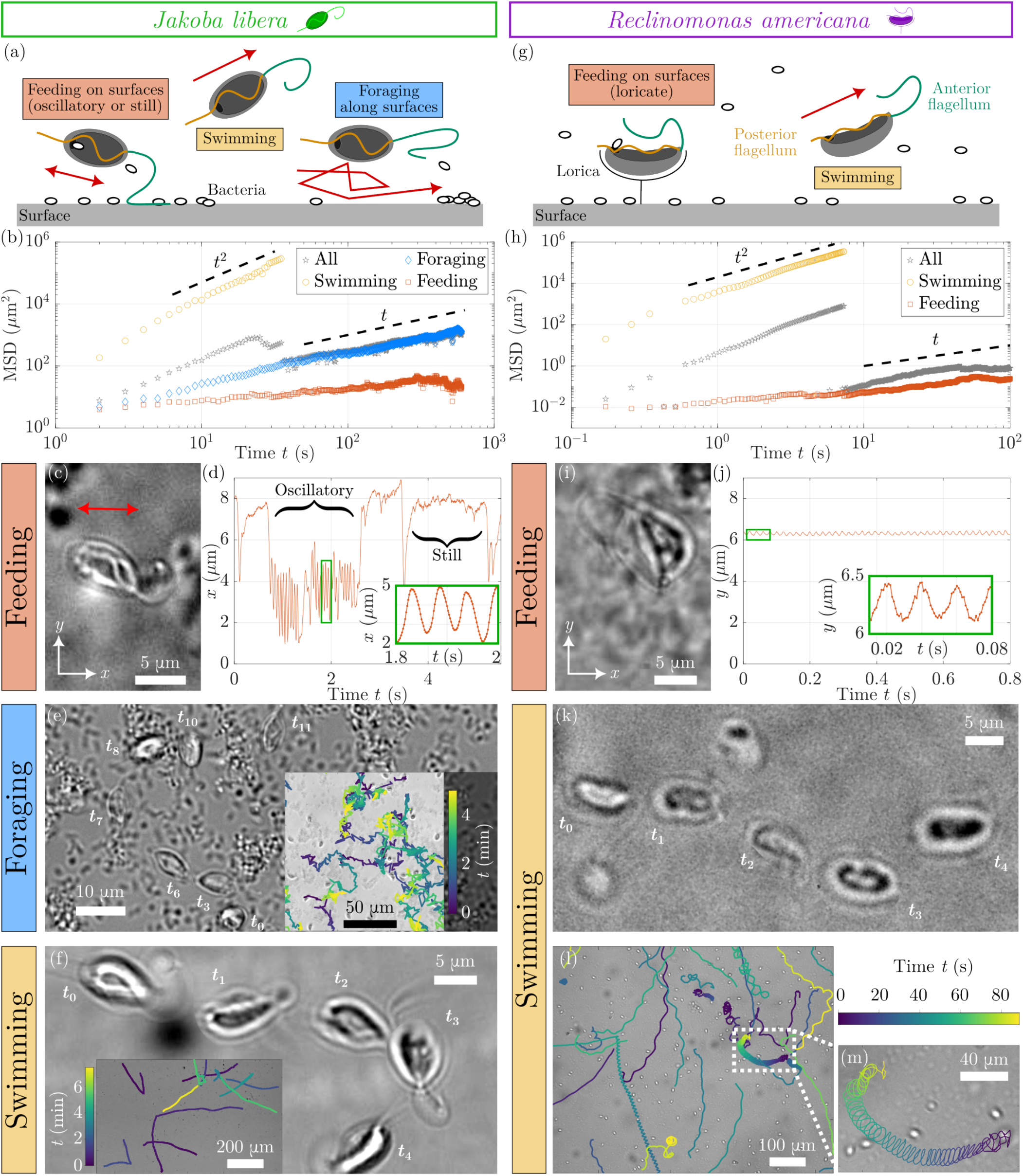
Comparison of *J. libera* and *R. americana* behaviours. (a,g) Illustration of the different behaviours. Red arrows indicate the direction of motion. (b,h) Mean squared displacement (MSD) as a function of time *t* for the entire population (grey pentagrams), as well as its sub-populations classified into swimming (yellow circles), feeding (orange squares), and foraging (blue diamonds, for *J. libera* only). (c-f) Three distinct behaviours in *J. libera*. (c) A feeding cell attached to a bacterial biofilm is imaged at 500 fps (see Movie S1). (d) Tracked *x* position of the same cell, exhibits both ‘oscillatory’ and ‘quiescent’ phases. (e) Chronophotography of a foraging cell on a surface (d*t* = 40 s), see Movie S2. Inset: typical trajectories. f) Chronophotography of a swimming cell in the bulk (d*t* = 1 s) (see Movie S3 and Fig. S3). Inset: typical trajectories. (i-m) Two distinct behaviours in *R. americana*. (i) A feeding cell is recorded at 1000 fps (see Movie S4). Tracked *y* position of the same cell shows its tethered state with small oscillations. (k) Chronophotography of a swimming swarmer cell in the bulk (d*t* = 0.14 s), see Movie S5. Inset: typical trajectories. (m) Zoomed-in view of a single helical trajectory with small pitch.

To observe single-cell feeding events, we used high-speed imaging (100 *-* 1000 fps) at 101× magnification (see Methods). A captured bacterium is typically guided along the ventral groove by the posterior flagellum, which adjusts its beating pattern to transport the prey to the posterior end for phagocytosis into the cytopharynx, where ingested bacteria accumulate [24]. Cell tracking reveals that *J. libera* alternates between oscillatory motion and a quiescent state, exhibiting minimal body movement during the latter (see Fig. 2c,d, also Movie S1). A cell can remain quiescent for extended periods, performing one or multiple oscillations between these periods. Oscillations have a period Δt of ≈ 70 ms (n = 5 cells) and an amplitude A = (1.6 *±* 0.4) µm (0.5 body size), see inset of Fig. 2d. We deduced that the oscillations result from the periodic beating of the anterior flagellum, which remains attached at one end to the substrate; the body movement is thus dominated by the activity of the anterior flagellum. The measured periodicity of body oscillations f = (14 *±* 3) Hz matches the beating frequency of the anterior flagellum (see also [10]). Meanwhile, the posterior flagellum beats consistently within the groove and contributes little to the body displacement.

The foraging behaviour of *J. libera* may be a previously undescribed form of microbial motility. The cells ‘jump’ or move fleetingly from one spot to another, searching suitable places to feed, and constantly attach and detach from the surrounding bacterial biofilm (see Movie S2 and Fig. 2e). Close-up trajectories of excavates exhibiting foraging are shown in Fig. 2e (inset). At long times, this motion produces a diffusive random walk in 2D along the surface (Fig. 2b). To better resolve the early-time characteristics of the foraging behaviour, sampling rate was increased to 100 fps. Here, the MSD displays (Fig. S4) two diffusive regimes at distinct timescales. We interpret the first as due to diffusion around a singular bacterial hotspot at very short times, and the second due to diffusion between different hotspots at longer times (see Movie S2). The first diffusive regime has a higher diffusion coefficient (*D* ≈ 5.9 μm^2^ s^−1^) than the second (*D* ≈ 0.75 μm^2^ s^−1^). While surprising, this higher value likely stems from the higher velocity (*v* ≃ 2*A* ⋅ 2*f* ≈ 50−140 μm s^−1^) of the excavate when it is attached, compared to when it is exploring the surface (*v* < 6 μm s^−1^). The two diffusive regimes are separated by a critical transition timescale of *t* = 0.2 − 3 s, corresponding to the residence time of the cells around one hotspot. In the context of optimal foraging theory, this adaptive behaviour could allow the excavates to efficiently scour the surface, switching from a more diffusive-like local search pattern when the density of prey is high, to a Lévy-like search pattern when prey density is more patchy [40, 41].

The bulk swimming behaviour of *J. libera* is predominantly due to the anterior flagellum which beats in a 3D

The bulk swimming behaviour of *J. libera* is predominantly due to the anterior flagellum which beats in a 3D motion away from the cell body to produce a characteristic back-and-forth movement of the cell. The anterior beat frequency of *f* ≈ 15 Hz is similar to that observed when feeding. The cell follows helicoidal trajectories, rotating on itself due to the asymmetrical 3D beating motion of the anterior flagellum, see Fig. 2f and Movie S3. The ‘rolling’ frequency is *f*roll ≈ 1.5 Hz, see Fig. S3, similar to other algal flagellates [17]. The posterior flagellum extends farther than the length of the cell body, but only the portion inside the groove beats vigorously in a 2D plane, while the distal portion beyond the cell is inactive. Transient changes in the anterior flagellum beat pattern reorient the cell before the usual waveform resumes (Fig. 2f and Movie S3). The motility of *K. bialata* is similar to that of *J. libera* (see Fig. S5), with cells swimming, foraging, or feeding (albeit the feeding behaviour is slightly different as *K. bialata* uses the posterior end of its body to attach to the surface instead of its anterior flagellum [10]). *R. americana* behaves differently. Loricate cells feed while attached to a surface (see Fig. 2i and Movie S4), by generating a feeding current with both flagella. The flagella movement also causes oscillations, see inset of Fig. 2j, of small amplitude *A* = (0.3 ± 0.1) μm (< 0.1 body size) and frequency *f* = (53 ± 4) Hz (*n* = 4 cells) − consistent with the anterior flagellum beat frequency reported for feeding *R. americana* [10]. No foraging behaviour is observed, unlike in the other two typical excavates. Swimming trajectories also have higher variability ranging from near straight paths to helices with small pitch, see Fig. 2k-m.

Next, we ask how these behaviours relate to the vertical distribution of excavates in the water column. From our holotomography images, we expect the excavates to be enriched near surfaces, this was confirmed and quantified by performing *z*-stack imaging from the bottom surface (*z* = 0) to the free surface at the top (see Methods). The concentration profiles of cells are plotted as a function of height (Fig. 3a,d). Most excavates are found within the first 100 μm. Above 100 μm, cell concentration decreases sharply, reaching a plateau of 2 − 4% of the maximum concentration *c*_0_ = *c*(*z* = 0) (concentration at the bottom surface) for *J. libera* and less than 0.1% for *R. americana*. These measurements show that most excavates are found close to the bottom surface, but with a small percentage of bulk-swimming cells. Surprisingly, the density at the free-surface (liquid-air interface) increases, reaching twice that in the bulk for *J. libera* (around 6 − 7% of *c*_0_), and nearly 100 times higher for *R. americana* (6 − 7%). This is confirmed by higher-magnification observations for *J. libera*, which reveal some excavates attached to a thin bacterial biofilm formed at the upper free-surface, where they perform the same feeding behaviour as observed on the bottom surface. This accumulation of bacteria at the free-surface could arise from aerotactic effects [15, 42, 43].

**Figure 3.**
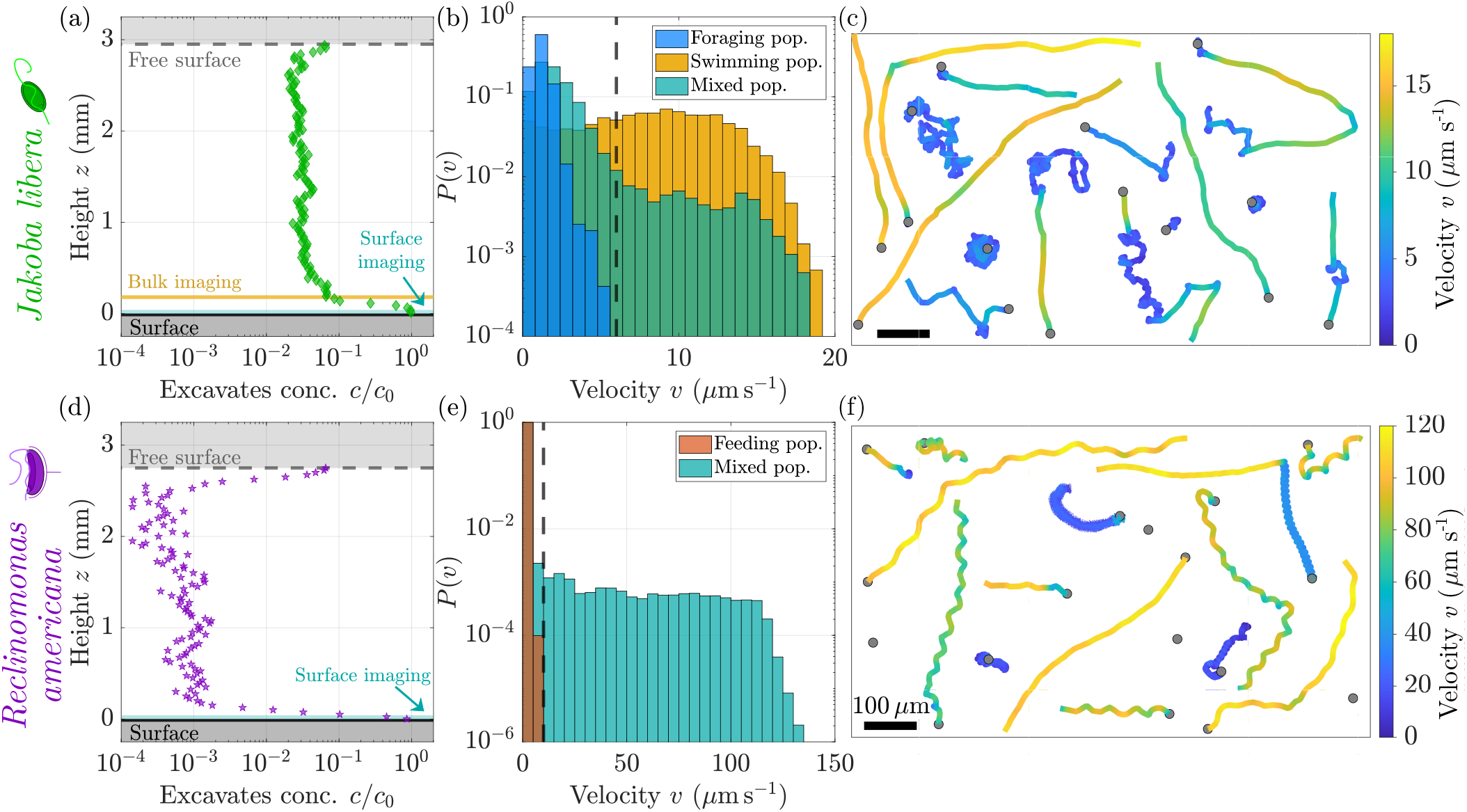
Motility transitions in *Jakoba libera* (a-c) and *Reclinomonas americana* (d-f). (a,d) Measured vertical concentration profile *c*(*z*) of *J. libera* (12-day old culture in ≈ 3 mm-deep suspension) and of *R. americana* (8-day old culture in ≈ 2.8 mm-deep suspension), both normalised by the concentration at the bottom surface *c*_0_ = *c*(*z* = 0). (Yellow and teal lines depict the depths of field of the 10× objective used.) (b,e) Probability distributions *P*(*v*) for representative *J. libera* and *R. americana* populations. *J. libera*: foraging-only (blue, day 3), swimming-only (yellow, day 13), or mixed (teal, day 14). *R. americana*: feeding-only (orange, day 14) or mixed (teal, day 7). Imaging was performed near the surface for the foraging-only and mixed populations (*z* = 0 − 40 μm), and in the bulk (*J. libera* only) for the swimming-only population (*z* = 160 − 200 μm). Vertical dashed lines: threshold used to separate swimming from the foraging (*v* = 6 μm s^−1^ for *J. libera*), and feeding cells (*v* = 10 μm s^−1^ for *R. americana*). (c,f) Typical trajectories of *J. libera* (14-day old culture) and *R. americana* near surfaces (7-day old culture), coloured by instantaneous velocity *v*. Grey circles mark the starting point of the tracks. Scalebars: 100 μm.

Motivated by these findings, we restrict ourselves hereafter to surface imaging and particularly behavioural characterisation at the bottom surface only. Custom velocity thresholds were used to distinguish between the swimming and the foraging or feeding behaviours. Fig. 3b,e show probability distributions of the instantaneous velocity *P*(*v*) for different populations (see Methods). Note that the maximum velocities measured for *R. americana* (120 μm s^−1^) are several times higher than for *J. libera*, which do not exceed 20 μm s^−1^ (see also Fig. 4), likely due to the much higher flagellar beat frequencies. Transitions from one motility mode to another also occur within a single trajectory (Fig. 3c,f).

**Figure 4.**
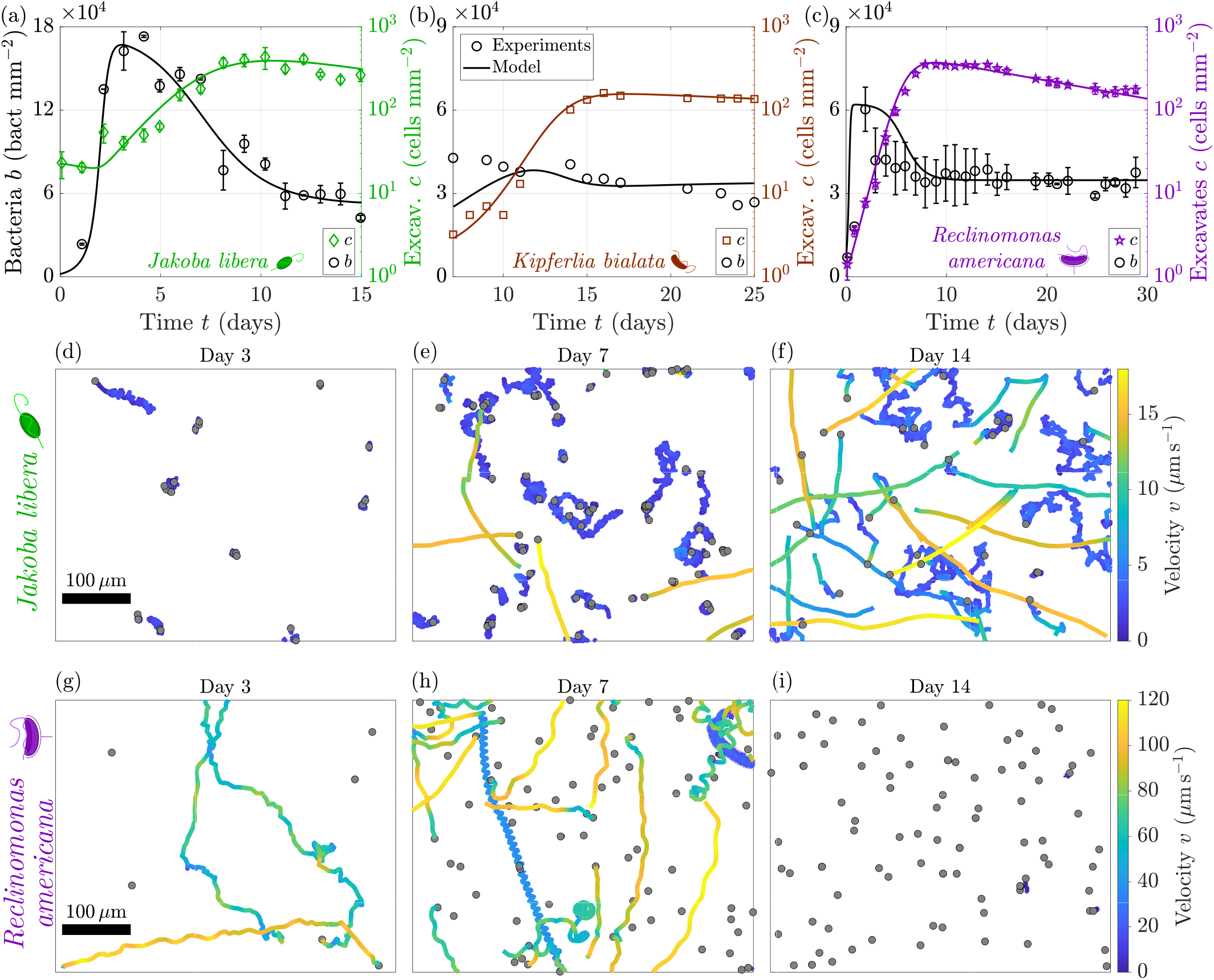
Growth and motility of typical excavates over time. (a-c) Growth curves of the excavates and bacteria for three species of typical excavates: (a) *Jakoba libera*, (b) *Kipferlia bialata*, and (c) *Reclinomonas americana*. Points: Experimental data, mean *±* SD*/*2 shown in (a,c) for two biological replicates. Lines: Fits to predator-prey model (*b* = *b*_*e*_ + *b*_*r*_, see Methods). (d-f) Trajectories of *J. libera* imaged over 10 min near the bottom surface, in (d) early growth-phase (*t* = 3 days), (e) late growth-phase (*t* = 7 days), or (f) saturation phase (*t* = 14 days), corresponding respectively to high, intermediate or low bacteria densities. (g-i) Trajectories of *R. americana* imaged over 1.5 min near the bottom surface, sampled at equivalent phases of the culture cycle as for *J. libera* above. Grey points mark the starting point of the tracks. (Fig. S5 for trajectories of *K. bialata*.)

### C. Dependence of motility behaviour on prey availability

For bacteriovorous flagellates, we anticipate the access to prey to be a natural driver of adaptive behaviours, as observed in other protists [44]. To test this, we conducted extensive behavioural imaging experiments over typical batch culture cycles (2 - 4 weeks depending on species), to quantify how these behaviours and their relative distributions change in the presence of environmental cues. Movies were recorded daily until saturation phase with a 10× objective, as well as snapshots of the surface with a 40× objective to measure the local bacteria density (see Methods). The growth curves of *J. libera* and their bacterial prey follow predator-prey dynamics, see Fig. 4a. A modified Lotka-Volterra model with four populations (predatory excavates, edible bacterial prey, resistant bacteria, and nutrients for bacteria) successfully reproduce the observed trends, see lines in Fig. 4a. See Methods and Table S1 for details on the model and the fitting parameters. The bacterial concentration b initially grows rapidly, before stabilising around 1.5 × 10^5^ bact mm^-2^ on day 3. After one week, b slowly decreases, reaching a density closer to that on day 1, i.e., 24 h after the beginning of the experiment. Meanwhile, the concentration c of the predator grows exponentially for 8 days, with a doubling time of 1.9 days, before reaching a plateau around 3 × 10^2^ cells mm^-2^. Similar growth curves of *K. bialata* and *R. americana* were obtained, see Fig. 4b-c, with exponential growth of the concentration of excavates followed by a plateau. The doubling times of *K. bialata* and *R. americana* are respectively 1.5 days and 0.8 days, and saturation concentrations of 1.5 × 10^2^-4 × 10^2^ cells mm^-2^. The model successfully reproduces the observed dynamics for these two species (fit parameters see Table S1).

Over time, the behaviour of the excavate predators changes dramatically. For *J. libera*, in the first 5 days all cells remain on the surface in the ‘feeding’ state with little motility, with a small proportion gently exploring the surface in ‘foraging’ mode (Fig. 4d). Lone swimming cells are seldom observed before day 6, with swimming trajectories only becoming noticeable after day 6, when the foraging cells also become more prevalent, see

Fig. 4e. As the culture ages, the motility of *J. libera* continues to increase steadily as quantified by the number of swimming cells, see Fig. 4f. *K. bialata* cultures exhibit a similar temporal dependence, see Fig. S5. While the motility of *J. libera* and *K. bialata* increases over time (similar to the choanoflagellate *Minosiga* [44]), *R. americana* exhibits the opposite behaviour. More specifically, the highly motile sub-population of *R. americana* diminishes when its concentration reaches saturation. This sub-population seems to exist only for as long as the number of excavates continues to increase, suggesting that cell division could be the only way for *R. americana* to become motile. Abundance of food could trigger cell division, one of the two daughter cells (the aloricate swarmer) would swim rapidly away from its loricate parent cell [30]. This parallels the asymmetric cell division and dispersal strategy of *Caulobacter* bacteria, while a stalked non-motile cell remains attached to the substrate, the motile swarmer cell swims away to colonise new habitats [45].

To characterise these trends at a population level, we systematically quantify the proportion of swimming cells over time, their swimming velocity, and diffusivity of the foraging state (where applicable). All imaging was performed at the bottom surface. We applied an instantaneous velocity threshold to all recorded trajectories (see Methods) to estimate c_*s*_ = c_swim_/c_0_, the proportion of swimming cells relative to the surface concentration c_0_ at z = 0 (c_0_ will be called c hereafter for simplicity). The time evolution of the swimming population is shown in Fig. 5a. On day 0, the proportion of swimming *J. libera* is around 3 %, but drops to a negligible amount (< 0.05 %) by day 2. Subsequently, this proportion increases again in a near-exponentially manner, reaching approximately 4 % by the end of the culture cycle. A higher percentage of swimming cells is also observed for *K. bialata* when food availability is low, although the increasing trend is not as clear due to limited statistics (see Methods). In contrast, *R. americana* showed a steadily decreasing trend of c_*s*_ as a function of culture age.

**Figure 5.**
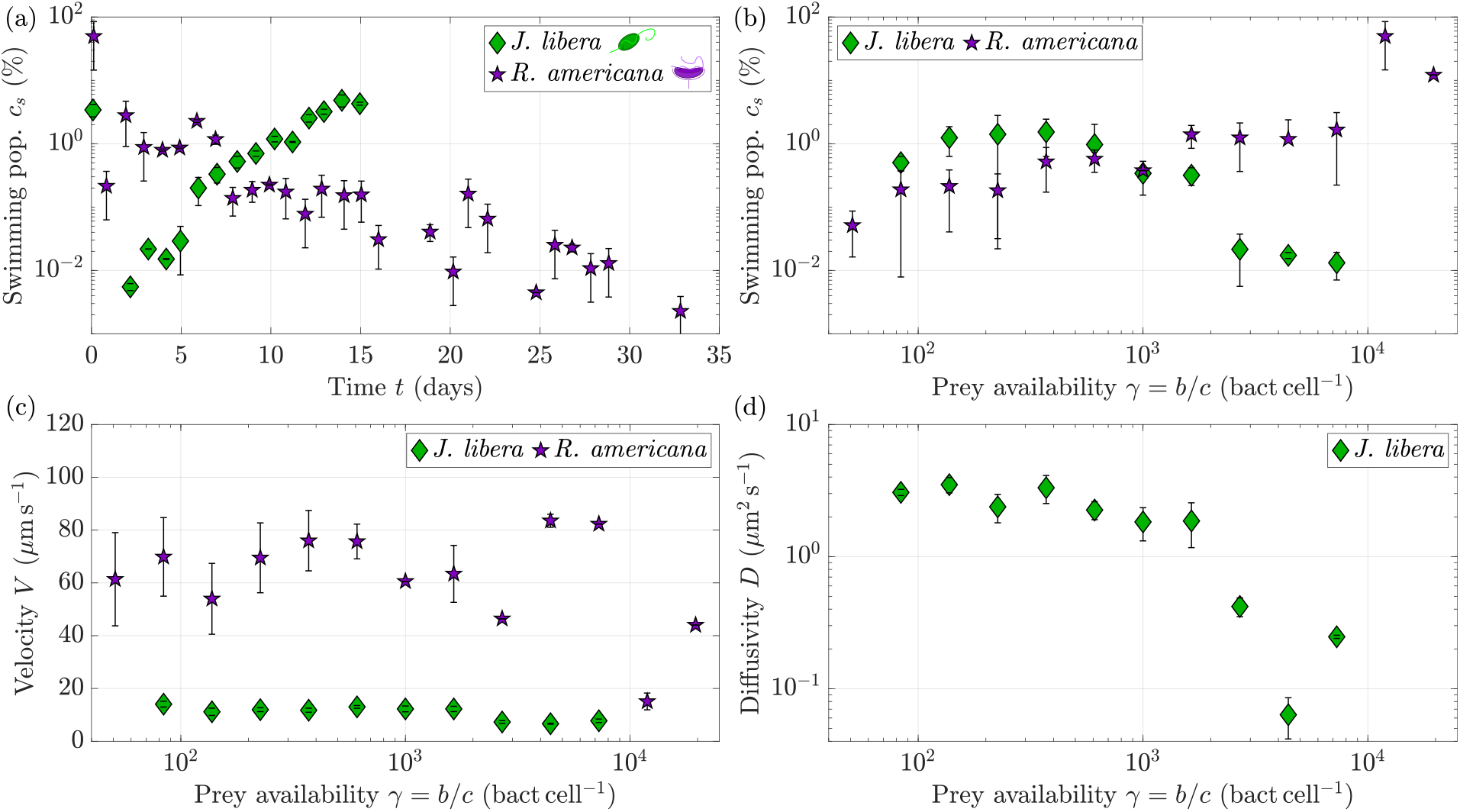
Motility of typical excavates varies with prey availability. (a) Temporal dynamics of the percentage of swimming excavates *cs*, for *Jakoba libera* (green diamonds) and *Reclinomonas americana* (purple pentagrams). Data points in (a) are from two biological replicates (see Methods), mean ± SD/2 is shown. (b) Percentage of swimming population *c*swim/*c* as a function of the ratio of the number of bacteria per excavate *γ* = *b*/*c*. (c) Mean velocity *V* of the swimming population, as a function of *γ* = *b*/*c*. (d) Diffusion coefficient of the foraging population of excavates, as a function of *γ* = *b*/*c*. Data points in (b-d): *n* = 100 experiments for *J. libera* (from 4+1 biological replicates), and *n* = 183 experiments for *R. americana* (from 4+2 biological replicates), see Methods. Filled symbols: logarithmic binning of the data in *γ*, mean ± SD/2 is shown.

Prey availability, which we define to be *γ* = *b*/*c*, the relative number of prey (bacteria) per predator (excavate), appears to be the important variable in regulating *J. libera* motility, rather than prey density itself. For example, according to the growth curves in Fig. 4a, *b* is similar on days 2 and 7 but the swimming population *cs* increases by two orders of magnitude (Fig. 5a). Fig. 5b correlates the observed adaptive behaviour with prey availability *γ* instead of time. For *J. libera*, the motility is low when the bacteria density is high (days 2 − 7), but high when the bacteria density is low (day 0 and days 8 − 15), leading to an increasing population of swimmers as food availability decreases up to a peak value at *γ* ≤ 800. In contrast, the proportion of swimming cells increases with *γ* for *R. americana*. Interestingly, the mean velocity *V* of the swimming cells was not affected in either species, suggesting that the adaptive behaviour mainly impacts the proportionality of the different behaviours. However, the diffusion coefficient *D* extracted from the foraging sub-population of *J. libera* cells exhibits a clear dependence on food availability *γ* (Fig. 5d). *D* drops by two orders of magnitude from ≈ 1 × 10^1^ μm^2^ s^−1^ to less than 1 × 10^−1^ μm^2^ s^−1^, when *γ* decreases from 10^4^ to 3 × 10^3^. This behaviour is consistent with our previous qualitative observations that foraging cells explored ever increasing distances when prey availability decreased (Fig. 4d-f). Similar trends are observed for *K. bialata* (see Fig. S5).

### D. Distribution and spatial statistics of surface-associated excavates

Next, we examine the distribution of excavates on surfaces, which we expect to depend on the adaptive motility and feeding strategy of each species. These distributions were quantified using spatial statistics tools to provide a population-level perspective on how predatory excavates colonise the surface. We focus on *J. libera* and *R. americana*. From the same videos used previously for the motility analysis, we first identified the centroid positions of the near-surface excavates on each frame (*z* = 0 − 40 μm). Voronoi tessellations were then constructed from these positions [46, 47] (see Methods). Examples are drawn in Fig. 6a-b for two experiments, for *J. libera* and *R. americana* respectively, when both have a similar surface density (*c* ≈ 35 cells mm^−2^).

**Figure 6.**
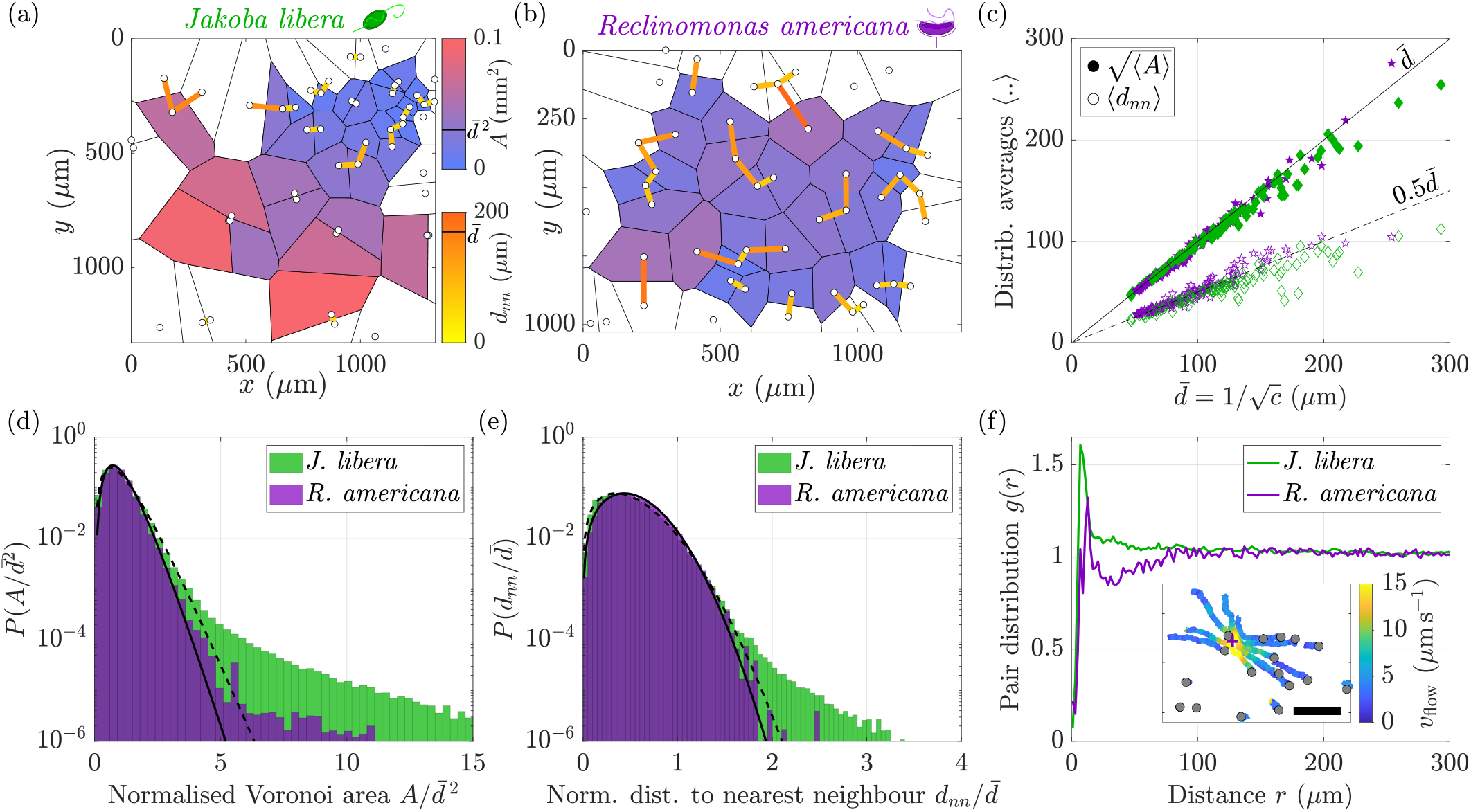
Spatial distribution of surface-associated excavates. (a,b) Voronoi tessellations constructed from excavate cell positions, for (a) *Jakoba libera* and (b) *Reclinomonas americana*, with surface imaging at 10× magnification (*z* = 0 − 40 μm). Closed Voronoi cells are coloured by area *A* from blue (small) to red (large), and nearest-neighbour distances *dnn* coloured from orange (small) to yellow (large). The same colourmaps are used for both species, since both are sampled at similar surface density (*c* ≈ 35 cells mm^−2^). (c) Ensemble averages of ⟨*dnn*⟩ and 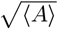 as a function of mean inter-excavate distance 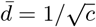. (d) Probability distributions 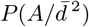, averaged across all experiments. 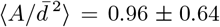 for *J. libera* and 0.95 ± 0.48 for *R. americana* (mean ± SD). Lines: fits of the distributions with gamma distributions (dashed for *J. libera* and solid for *R. americana*). (e) Probability distributions 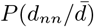, averaged across all experiments. 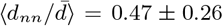 for *J. libera* and 0.51 ± 0.25 for *R. americana* (mean ± SD). Lines: fits of the distributions with Weibull distributions (dashed for *J. libera* and solid for *R. americana*). (f) Pair distribution function *g*(*r*) averaged across all experiments. Inset: Particle trajectories of sessile bacteria around a loricate *R. americana*, used as proxy for the feeding flow field *v*flow (see Fig. S9 and Movie S7). Purple cross: excavate location. Scalebar: 50 μm.

There is a clear difference between the two tessellations, which we quantified using the Voronoi polygon area A and the nearest neighbour distance d_*nn*_ (Fig. 6a-b). Whereas Voronoi polygons are similar in size for *R. americana*, their sizes vary greatly for *J. libera*. Indeed, foraging *J. libera* can form small clusters of high cell density around patches or hotspots of bacteria, meaning that the simultaneous existence of dense and empty regions is likely, unlike *R. americana* which creates long-range feeding flows (see Fig. S9 and Movie S7). This also explains the greater variance in the distances d_*nn*_ observed for *J. libera* compared to *R. americana*.

To quantify population-level trends, we measured *A* and *d*_*nn*_ on every frame of every motility video recorded (*n* = 100 videos for *J. libera* and *n* = 183 for *R. americana*, 600 or 1200 frames per video, see Methods). We then compared the mean inter-excavate distance 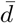 (estimated from the surface density *c* of the excavates via the relation 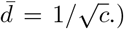, with the ensemble averages 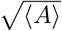 and ⟨*d*_*nn*_⟩ across all frames of each video, see Fig. 6c. As expected, both 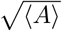 and ⟨*d*_*nn*_⟩ scale linearly with 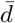, with 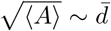 and 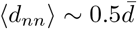.

Next, we obtain the distributions of Voronoi polygon areas and nearest neighbour distances, normalising *A* by 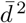 and *d*_*nn*_ by 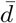. This allows us to aggregate data from experiments with different concentrations *c* of excavates. From a total of *n* = 1.1 × 10^*7*^ *−* 1.3 × 10^*7*^ data points for *J. libera* and *n* = 3.6 × 10^*7*^ *−* 4.0 × 10^*7*^ for *R. americana*, we obtained distributions with probabilities resolved down to 1 × 10^*−6*^ (Fig. 6d-e). These distributions confirm what is observed qualitatively in the snapshots above. Both distributions show a similar overall trend for the two species, but with some differences in the tail regions at the more extreme values. Compared to *R. americana, J. libera* distributions have longer tails. This difference arises from the slight clustering of excavates around bacterial hotspots which increases both the small distances between them and the probability of finding ‘empty’ regions with few cells. For *R. americana*, the normalised Voronoi areas follow a gamma distribution (shape factor *α* = 3.89 and scale factor *θ* = 0.25), while the normalised distances to nearest neighbour follow a Weibull distribution with a shape factor *k* = 2.16 close to 2 (and a scale factor *λ* = 0.58 close to the expected value 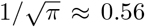 for a normalised dataset), see Supplementary Material for their mathematical expressions. These distributions are consistent with a homogeneous Poisson process in 2D, where cells are distributed randomly across a surface [48, 49]. For *J. libera*, the central region of both distributions is also well characterised by, respectively, a gamma distribution (*α* = 2.79 and *θ* = 0.34) and a Weibull distribution with a shape factor close to 2 (*k* = 1.87 and *λ* = 0.53), but diverges from these fits at very low and very high values.

To further assess spatial variations, we measured, using the same data, the pair distribution function *g*(*r*) of the excavates on the surface, defined as the probability of finding a pair of particles separated by a distance *r* (see Methods), see Fig. 6f. For a perfectly homogeneous Poisson process, *g*(*r*) = 1. Here, both species *g*(*r*) deviate from this baseline for small values of *r*. After a region of exclusion below one cell width (presumably due to steric effects), pronounced peaks are observed. For *J. libera*, this peak is located at *r* ≃ 7 μm (~ body size), indicative of short-range structure, likely due to clustering around bacterial hotspots. Then, *g*(*r*) slowly decreases to 1, meaning that the distribution tends to uniformity. For *R. americana*, the initial exclusion region is slightly larger, likely due to the lorica. A small peak is again observed at *r* ≃ 7 − 13 μm, followed by a zone of weak repulsion, that reaches a minimum at *r* ≃ 30 − 35 μm and extends until *r* ≃ 75 μm. This is similar to the range of hydrodynamic interactions generated by feeding flows of attached cells (see Fig. S9 and Movie S7).

Finally, we explore how these spatial statistics depend on the density *b* of bacteria, *c* of excavates, or prey availability *γ* = *b*/*c*, by binning the data according to measurements of these parameters made separately on the same video microscopy dataset, and recalculating the distributions of *A, dnn*, and *g*(*r*). The scaling with respect to *c* was the most informative, see Fig S10a-i. The long tails of the distributions of *A* and *dnn* for *J. libera* appear predominantly at high *c*, while the short-range clustering is enhanced at low *c*. For *R. americana*, no noticeable differences are visible with *c*. To capture density fluctuations over larger length scales, we also computed the structure factor *S*(*k*) (Fig S10j-l) and number variance *σN* (Fig. S11), which can also be used to assess hyperuniformity. To account for finite size effects, we compared the results with Poisson simulations in equivalently sized domains (see Methods). *S*(*k*) and *σN* differ between the two excavates, and from the Poisson simulations. Deviations from the expected Poisson scalings were observed with weak dependence on the excavate cell density *c*, including enhanced long-range interactions in *J. libera* contrasting with suppressed fluctuations (hyperuniformity [32, 50]) in *R. americana*, as *c* increases (Fig S11).

## DISCUSSION

### Prey-dependent adaptive behaviours

In this article, we have identified and compared the adaptive motility and feeding behaviours of three excavate protists: *Jakoba libera, Kipferlia bialata*, and *Reclinomonas americana*, showcasing a multiscale approach that is applicable to a wide range of organisms. We extensively quantified excavate behaviours in typical batch cultures over successive days. Important differences were revealed across species, particularly in how cells select between behavioural states (a form of decision-making). We identified a major driver of this selection to be the availability of their co-cultured bacterial prey (see Fig. 4). So far, behavioural changes with food or prey abundance have only been sparsely reported [28, 44, 51–53], with some larger species exhibiting rich collective behaviour [54] or even phenotypic transitions [55, 56]. While most laboratory studies focus on short-time observations and qualitative descriptions or measures of growth kinetics, the present study provides a comprehensive picture of how the behaviour of microbial hunters dynamically shifts in response to changing prey abundance.

Our analysis of these interactions showed predator-prey dynamics in the excavate-bacteria densities, with strong dependence of motility metrics on *γ* = *b*/*c*, the number of bacteria available per excavate (see Fig. 5). For *J. libera* and *K. bialata*, motility increases with decreasing *γ*, whereas the opposite trend is observed for *R. americana* which becomes more likely to attach to surfaces in its loricate form. When food is limited, *J. libera* expands its search radius by modulating the diffusion coefficient of foraging by ~ 2 orders of magnitude, or by increasing the proportion of swimming cells (also by 2 orders of magnitude, 5). By dynamically adjusting their motility, *J. libera* and *K. bialata* actively seeks out new habitats when the current one becomes exhausted, while *R. americana* relies predominantly on effective dispersal after cell division by fast-swimming swarmer cells to colonise new habitats. The strong dependence of the motility on prey availability, rather than prey density, hints at possible competition between predators. Cells may be vying to occupy the same bacterial hotspots (see Movie S6 and Fig. S8).

### Morphology and flagellar control of behaviour

The species share a strikingly similar morphology, which we quantified using U-ExM, producing detailed measurements of cell and groove shape, see Fig. 1, Fig. S1 and Table. S2. High-speed imaging helped reveal how flagellar dynamics contribute to interspecies differences in behaviour, as well as shifts in behaviour within the same individual. Three characteristic behaviours were identified for *J. libera* and *K. bialata*: feeding on surfaces, foraging along surfaces, and swimming in bulk fluid, see Fig. 2. Near surfaces, *J. libera* alternates between a ‘quiescent’ and an ‘oscillatory’ state dominated by the regular beating of the cell’s anterior flagellum, which remains attached to the surface. Foraging involves two distinct diffusive timescales, corresponding to diffusion around a singular bacterial hotspot at short times, and between hotspots at longer times. For *R. americana*, foraging behaviour is not observed and their feeding behaviour only consists of a ‘quiescent’ phase where the cell remains attached to the surface by its lorica while both flagella beat to generate a strong feeding current.

Motile microbes often exhibit variable behaviours including stochastic transitions between motility states [38, 39, 57, 58], particularly near boundaries, surfaces, or in confinement [37, 59]. In excavates, behavioural variability was evident in single-cell trajectories not just as a population-level phenomenon, but also in the spontaneous and reversible transitions between foraging and swimming in *J. libera* (Fig. 3). Our observations show that different behavioural modes are readily accessed by modulating the dynamics of one or both flagella, enriching our perspective on the option space of morphology-function relationships in simple flagellates [60].

### Ecological significance of surfaces

Motility has clear ecological consequences for competition in spatially-distributed resources [61]. Under the heterogeneous, time-varying environmental conditions inhabited by microbes, excavate predators must balance the exploration of new territory with the exploitation of local resources. Our work highlights the importance of surfaces on the ecology and behaviour of typical excavates [10, 24]. Measurements of the vertical concentration profiles within the samples show that excavates overwhelmingly occupy the bottom surface, with a small percentage at the free surface, and very few in bulk fluid far from surfaces. Accumulation at the free surface was reported in other eukaryotes [62–64], indicating this behaviour may be universal. Paramecia also immobilise near surfaces by reducing their cilia activity, depending on the availability of bacteria [65]. Here, we measured the consequences of different adaptive motility and feeding behaviours on the time-varying spatial distribution of excavates on surfaces through rigorous tracking and statistical quantification (Fig. 6), again highlighting interspecies differences. These patterns may facilitate collective resource acquisition, similar to the distribution of photoreceptor cells in the avian retina [66] or dinoflagellates at air-liquid interfaces [50].

### Evolutionary perspective

Given their presumed morphological and phylogenetic proximity to LECA [26, 27], the finding of behavioural plasticity in modern-day excavates suggests that these traits may have already been present in LECA. The capacity for adaptive foraging strategies may have given these deep-branching protists a considerable selection advantage during the diversification of eukaryotes. The findings raise the important question of how protistan predators perceive these environmental changes (such as prey abundance), and how this information is transduced to the ciliary motor apparatus to dictate the cell’s motility state. The integration of sensory processing, motor control, and decision-making in a single-celled organism constitutes a minimal form of cellular cognition. Future work should also seek to determine how dynamic predator-prey interactions could drive changes in community composition across ecologically relevant timescales.

## MATERIALS AND METHODS

### Culture preparation

The typical excavates *Jakoba libera* (ATCC 50422), *Kipferlia bialata* (WC1A) and *Reclinomonas americana* (BTAS1C) were kindly provided by Alastair Simpson (Dalhousie University) and Sei Suzuki-Tellier (Technical University of Denmark). All cultures were kept in the dark at ambient temperature (≈ 20°C). *J. libera* and *R. americana* were subcultured into horizontal culture flasks every two (*J. libera*) or four weeks (*R. americana*), while *K. bialata* was cultured in vertical 15 mL Falcon tubes every week in anoxic conditions. The marine species *J. libera* and *K. bialata* were grown in Artificial Sea Water medium, prepared using TMC Premium Reef-salt at 30 parts per thousand. The medium is supplemented with, respectively, 0.3% and 3% of pure LB (Luria-Bertani), a rich medium providing the necessary nutrients to the bacteria the excavates feed on. *R. americana*, a fresh-water species, was grown in milli-Q water with 0.3% of pure LB.

### Experimental setups & Imaging

#### Motility experiments

A 1 mL sample was placed into an Ibidi 2-well μ-Slide with a glass coverslip bottom (Ibidi, Germany). The μ-Slide was then positioned on either an Olympus IX83 inverted microscope (Olympus, Japan) or a Leica DMI8 inverted microscope (Leica Microsystems, Germany), and the focussing was done on the lower surface of the specimen. Initially, a single movie is captured in bright-field using a 10× objective (depth of field ≈ 40 μm). These movies will be used to estimate the number of excavates and to extract their trajectories. We then switch to a 40× objective to capture an image, at the same location, which will be used to determine bacterial density. Movies were recorded with a 10× objective at 1 or 2 fps for *J. libera*, 2 fps for *K. bialata*, and 13.5 ± 0.2 fps for *R. americana*, using a pco.panda 4.2 camera (Excelitas, USA) or a Leica DMC2900 camera (Leica Microsystems, Germany). 10 min movies were recorded for *J. libera* and *K. bialata* (along some 20 s movies for *K. bialata*), and ≈ 1.5 min-long movies for *R. americana*. A faster acquisition rate was needed for *R. americana* due to its higher swimming motility.

#### Growth experiments

Growth experiments were conducted in an Ibidi 2-well µ-Slide for all three typical excavates. The slides were hermetically closed with a lid and some Parafilm® to avoid evaporation and contamination. The experiments started (day 0) with a sample from a freshly prepared passage. The growth of bacteria and excavates and the motility of the excavates were recorded, as previously described (see Motility experiments), every day for 2 to 3 weeks for *J. libera* and 4 to 5 weeks for *R. americana*. 4 biological replicates were done for *J. libera* and *R. americana*: 2 replicates were prepared from the same mother culture at 2 different times of the year. For *K. bialata*, experiments were conducted approximately every couple of days with 1 biological replicate for 4 weeks because of difficulties while culturing. Experiments without LB in the passage were also conducted for *J. libera* (1 replicate) and *R. americana* (2 replicates), showing nearly no bacterial growth and reduced excavate growth. Changes in the excavates motility over time were similar, but less significant. Data from these experiments is also included in Fig. 5b-d and Fig. 6c-f.

#### z-profiles

The excavates concentration profiles in *z* were performed in bright-field on an Olympus IX83 inverted microscope (Olympus, Japan) with a 01-Kinetix-M-C camera (Teledyne Photometrics, USA) and a 10× objective. *z*-profiles were taken in a 1 mL suspension with d*z* = 10 μm steps across the entire liquid height, in an Ibidi 2-well (up to 1 mm) or 8-well (up to 3 mm) μ-Slide.

#### High-speed imaging

A Phantom v1212 camera (Phantom, USA) on a Leica DMI8 inverted microscope (Leica Microsystems, Germany) was used for high-speed imaging. Movies of feeding excavates on surfaces were recorded at 100, 500 or 1000 fps (depending on the species) in bright-field with a 63× water immersion objective for individual cells, sometimes further magnified 1.6 times with a magnification changer (i.e. 101×). High-speed movies of foraging *J. libera* along surfaces were recorded at 100 fps with a 20× objective. Imaging was performed on 1-week-old samples (1 mL) in an Ibidi 2-well μ-Slide, originated from a newly prepared passage.

#### Expansion microscopy

Ultrastructure Expansion Microscopy (U-ExM) images were recorded with a 40× water immersion objective on a Zeiss confocal microscope with Airyscan (Zeiss LSM 880, Germany). The *z*-stacks were spaced by d*z* = 0.2 − 0.5 μm. Protocol followed Ref. [35]. Primary antibodies against *α* and *β*-tubulin (AA345 & AA344 guinea pig from ABCD Antibodies, Switzerland) were used in conjunction with the appropriate secondary antibody (Alexa Fluor− 568, A11075 from Thermo Fischer, Germany) to visualise tubulin. DAPI 4’,6-diamidino-2-phenylindole (D1306 from Thermo Fischer, Germany) was used for DNA visualisation. The expansion factor of the cells was estimated by measuring the expansion factor of the gel in which they are embedded.

#### Holotomography

Holotomography images were obtained with a Tomocube HT-X1− (Tomocube, Korea) on 3-week-old samples of *K. bialata* and *J. libera* in an Ibidi 2-well μ-Slide. The 3D stacks consist of (310 × 310) μm^2^ images, spaced by d*z* = 0.84 μm and centred on the bottom surface. The liquid medium surrounding the microorganisms was chosen as the reference for the refractive index mapping, with a refractive index identical to that of water, i.e. 1.3337.

### Data analysis

Data analysis was conducted with MATLAB (The MathWorks Inc.). Fiji [67] and TrackMate plugin [68] were used for cell detection and tracking. 3D reconstruction of holo-tomography and U-ExM images was made in DragonFly 3D World (Comet Technologies Canada Inc.). See Supplementary Material for more details.

#### Trajectories

For the growth experiments, the estimation of the number of bacteria was done by detecting circular spots of size ~ 1 μm in Fiji with TrackMate. The bacterial density *b* is thus a measure of the surface occupied by the bacteria (a long filamentous bacterium is not counted as one, but as multiple bacteria). For the movies recorded in 10×, the instantaneous velocity *v* of *J. libera* was smoothed with a moving average over a window of 5 s (see Fig. S3). The mean squared displacement (MSD) is defined as:

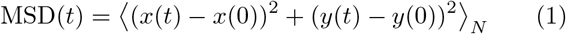

with N the number of excavate trajectories considered. We compute both the MSD of the entire population and of each sub-population using the criteria detailed in the Supplementary Material. The diffusion coefficient D is calculated from the MSD where relevant using the equation MSD = 4 *Dt* in an appropriate time range, since the recorded trajectories are in two dimensions.

To more accurately count the swimming population, a threshold of 6-10 µm s^-1^ (depending on the species and the biological replicate) is applied on the instantaneous velocity v. This allows us to take into account excavates transitioning from one motility mode to another.

#### Spatial statistics

For each image from every motility movie (n = 100 movies for *J. libera* and n = 183 for *R. americana*), spatial statistics are calculated on the positions of the excavates. Voronoi tessellations are constructed from these positions using Delaunay triangulation with the familiar Euclidean distance. Each Voronoi polygon contains only one excavate and represents the region of the plane closest to this specific excavate than any other [46, 47]. The Voronoi polygon area A and the nearest neighbour distance d_*nn*_ are used to quantify these tessellations (see Supplementary Material for their definition).

Three other metrics are used to quantify the surface distribution of excavates. The pair distribution function g(r) is the probability of finding a pair of excavates separated by a distance r. The structure factor S(k) measures their spatial organisation, in Fourier space, across different length scales. The number variance cr_*N*_ (R) is the variance in the number of excavates contained within circular regions of radius *R*. It is calculated by placing 5000 randomly positioned circles of radius *R* inside the observation area, and measuring how much the number of excavates inside these circles fluctuates. See Supplementary Material for their definition.

To identify whether trends observed in the spatial statistics characterisation are due to the experimental system or to finite-size effects, numerical simulations of a homogeneous spatial Poisson process are conducted in MATLAB. The simulations consist in generating random point patterns with a prescribed average density, inside a rectangular observation window identical to the experimental field of view. Ten average densities are prescribed, spanning the range of experimental excavate density c measured. 200 realisations are performed for each density. Results are shown in Fig. S10-S11.

A homogeneous spatial Poisson process is characterised by *g*(*r*) = 1, *S*(k) = 1 and a number variance cr_*N*_ scaling as *R*^2^. The distribution of its Voronoi areas A follows a gamma distribution, while the distribution of nearest neighbour distance d_*nn*_ follows a Weibull distribution with a shape factor k of 2 [48, 49]. See Supplementary Material for their definition.

#### Predator-prey model

To model the predator-prey dynamics, we use a modified version of Lotka-Volterra equations. We introduce a population of resistant bacteria not consumed by excavates (*b*_*r*_) and a population of nutrients (*n*_*u*_), alongside the prey (edible bacteria consumed by excavates *b*_*e*_) and the predators (excavates *c*). The assumptions are that nutrients *n*^*u*^ are consumed by both *b*_*e*_ and *b*_*r*_, the edible bacteria population *b*_*e*_ is consumed by excavates, and the excavates population *c* dies of starvation at a fixed rate. Each sub-population of excavates (feeding, swimming, and foraging where relevant) is assumed to behave similarly. See Supplementary Material for the set of differential equations describing the dynamics of the populations and the parameters used to fit the experimental data.

## ACKNOWLEDGMENTS

We thank Alastair Simpson, Sei Suzuki-Tellier and Thomas Kiørboe for fruitful discussions and for kindly providing the strains. We also thank Hannah Laeverenz-Schlogelhofer, Armando Rubio Ramos, Paul Guichard and Virginie Hamel for help with the expansion microscopy protocol, Bradley Dyer for help with the holotomography and expansion imaging, and Daryna Zavadska for sharing insights about protist behaviours.

This work is supported by the Human Frontier Science Program (Ref.-No: RGP014/2024) and the European Research Council (ERC) under the European Union’s Horizon 2020 research and innovation programme grant 853560 EvoMotion. This work also benefitted from state aid under France 2030 (Ref.: ANR-11-IDEX-003), and BBSRC internationalisation funds (BB/Y514147/1).

For the purpose of open access, the author has applied a Creative Commons Attribution (CC BY) licence to any Author Accepted Manuscript version arising from this submission.

## DATA AVAILABILITY

Data needed to reproduce all results in the paper is available at [insert URL after peer-review].

## CONFLICT OF INTEREST

The authors declare no conflict of interest.

## ARTIFICIAL INTELLIGENCE (AI) DISCLOSURE

AI was used for the creation of algorithms for data analysis for surface distribution analysis (ChatGPT-5.5), and proofreading and editing (Writefull).

## AUTHOR CONTRIBUTIONS

J.B. and K.W. conceptualised the research and designed the experiments. J.B. performed U-ExM experiments. J.B. and A.G. performed growth, motility and holotomography experiments. J.B. and S.B performed high-speed imaging experiments. J.B. performed simulations. H.A., A.G., and J.B. developed the predator-prey model. J.B. developed the methodology, analysed and visualised the data. K.W., J.B., and H.A. supervised the project and secured funding. J.B. and K.W. wrote the original draft of the manuscript. All authors discussed the results and reviewed the manuscript.

## Notes

### Competing Interest Statement

The authors have declared no competing interest.

