## Supplementary Information for "Adaptive motility and feeding strategies of deep-branching predatory protists"

Julien Bouvard,[1,*] Angéline Girard,[1,2] Samuel A. Bentley,[1] Harold Auradou,[2] and Kirsty Y. Wan[1,†]
[1] *Living Systems Institute, University of Exeter, Exeter, EX4 4QD, United Kingdom*
[2] *Université Paris-Saclay, CNRS, FAST, 91405 Orsay, France*
(Dated: October 7, 2026)

### I Data analysis

Data analysis was conducted with MATLAB (The MathWorks Inc.). Fiji [1] and TrackMate plugin [2] were used for cell detection and tracking. 3D reconstruction of holotomography and U-ExM images was made in DragonFly 3D World (Comet Technologies Canada Inc.).

#### A. Trajectories

For the growth experiments, the estimation of the number of bacteria was done by detecting circular spots of size $\approx 1\,\mu$m in Fiji with TrackMate. The bacterial density $b$ is thus a measure of the surface occupied by the bacteria (a long filamentous bacterium is not counted as one, but as multiple bacteria). The number of excavates was calculated from the movies recorded in 10× as the mean number of excavates throughout the entire movie, after having removed any outliers. For these movies, the instantaneous velocity $v$ of *Jakoba libera* was smoothed over a window of 5 s to reduce the noise originating from the helical motion of the excavates (see Fig. S3):

$$v(t) = \frac{1}{5} \sum_{\tau=t-2}^{t+2} \sqrt{v_x(\tau)^2 + v_y(\tau)^2} \qquad \text{(S1)}$$

with $v_x(t) = \frac{x(t+\mathrm{d}t)-x(t-\mathrm{d}t)}{2\mathrm{d}t}$ and $v_y(t) = \frac{y(t+\mathrm{d}t)-y(t-\mathrm{d}t)}{2\mathrm{d}t}$.

The mean squared displacement (MSD) is defined as:

$$\mathrm{MSD}(t) = \left\langle (x(t)-x(0))^2 + (y(t)-y(0))^2 \right\rangle_N \qquad \text{(S2)}$$

with $N$ the number of excavate trajectories considered. We compute both the MSD of the entire population and of each sub-population using the following criteria. For *J. libera*, a trajectory is considered 'swimming' if its median velocity $v_{\mathrm{med}}$ is higher than 6 or $9\,\mu\mathrm{m\,s^{-1}}$ (depending on the biological replicate); 'feeding' when the distance between the start and end points of its trajectory $d_{\mathrm{travel}}$ is shorter than $8\,\mu$m ($\sim$ 1 body size); and 'foraging' if both $d_{\mathrm{travel}} > 8\,\mu$m and $\max(v) < 6$ or $9\,\mu\mathrm{m\,s^{-1}}$. For *Kipferlia bialata*, the threshold on $d_{\mathrm{travel}}$ is identical to *J. libera* ($8\,\mu$m), while the threshold on the velocities $v_{\mathrm{med}}$ and $\max(v)$ is $9\,\mu\mathrm{m\,s^{-1}}$. For *Reclinomonas americana*, a trajectory is considered 'swimming' if its median velocity $v_{\mathrm{med}}$ is higher than $10\,\mu\mathrm{m\,s^{-1}}$, and 'feeding' when the distance between the start and end points of its trajectory $d_{\mathrm{travel}}$ is shorter than $4\,\mu$m ($\sim 1/2$ body size). Note that this categorisation is accurate but does result in some trajectories getting discarded, see Fig. S6. Most discarded trajectories exhibit transitions between one motility mode and another. The diffusion coefficient $D$ is calculated from the MSD where relevant using the equation $\mathrm{MSD} = 4Dt$ in an appropriate time range, since the recorded trajectories are in two dimensions.

To more accurately count the swimming population, a threshold of 6-10 $\mu\mathrm{m\,s^{-1}}$ (depending on the species and the biological replicate) is applied on the instantaneous velocity $v$. This allows us to take into account excavates transitioning from one motility mode to another, see Fig. S7.

#### B. Spatial statistics

For each image (600 or 1200) from every motility movie ($n = 100$ movies for *J. libera* and $n = 183$ for *R. americana*), spatial statistics are calculated on the positions of the excavates. Voronoi tessellations are constructed from these positions using Delaunay triangulation with the familiar Euclidean distance. Each Voronoi polygon contains only one excavate and represents the region of the plane closest to this specific excavate than any other [3, 4]. Note that only complete Voronoi polygons, *i.e.*, polygons fully contained in the field of view, are considered in our subsequent analysis. The Voronoi polygon area $A$ and the nearest neighbour distance $d_{nn}$ are used to quantify these tessellations. Voronoi polygons intersecting with the boundary of the observation window are excluded. For a given excavate $i$, its nearest neighbour distance is $d^i_{nn} = \{\min_{\forall j\neq i} d^{ij} \mid \min_{\forall j\neq i} d^{ij} < d^i_{\mathrm{edge}}\}$, with $d^{ij}$ the distance between two excavates $i$ and $j$, and $d^i_{\mathrm{edge}}$ the distance between the excavate $i$ and the closest edge of the observation window.

The pair distribution function $g(r)$ is the probability of finding a pair of excavates separated by a distance $r$. For an excavate density $c = N/A_{\mathrm{window}}$, with $N$ the total number of excavates and $A_{\mathrm{window}}$ the area of the

* Contact author:
† Contact author:

observation window, it is defined as:

$$g(r) = \frac{1}{cN} \sum_{i=1}^{N} \frac{N^i(r)}{A^i_{\text{accessible}}(r)} \tag{S3}$$

with $N^i(r)$ the number of excavates $j$ located at a distance $r \pm \Delta r$ from excavate $i$ (i.e., $r - \Delta r/2 \leq d^{ij} < r + \Delta r/2$) with $\Delta r = 2\,\mu\text{m}$, and $A^i_{\text{accessible}}(r)$ the accessible area of the annulus at a distance $r$ around excavate $i$ to account for the boundary of the observation window.

The structure factor $S(k)$ is defined as:

$$S(k) = \frac{1}{N} \left| \sum_{j=1}^{N} e^{-i(k_x x_j + k_y y_j)} \right|^2 \tag{S4}$$

with $N$ the number of excavates, $(x_j, y_j)$ the position of the excavate $j$, $k_x = 2\pi n_x/x_{\max}$ and $k_y = 2\pi n_y/y_{\max}$ physical wavevector components, $(x_{\max}, y_{\max})$ the size of the observation window, and $n_x$ and $n_y$ dimensionless integer Fourier-mode indices.

The number variance $\sigma_N(R)$ is the variance in the number of excavates contained within circular regions of radius $R$. It is calculated by placing 5000 randomly positioned circles of radius $R$ inside the observation area, and measuring how much the number of excavates inside these circles fluctuates. Its expression is:

$$\sigma_N(R) = \frac{1}{5000} \sum_{\alpha=1}^{5000} (N_\alpha(R) - \langle N(R) \rangle)^2 \tag{S5}$$

with $N_\alpha(R)$ the number of excavates in the $\alpha$-th circle and $\langle N(R) \rangle = (1/5000) \sum_{\alpha=1}^{5000} N_\alpha(R)$ the mean number of points. $R$ ranges from 1 to half of the smaller side of the observation window ($\approx 500\,\mu\text{m}$).

To identify whether trends observed in the spatial statistics characterisation are due to the experimental system or to finite-size effects, numerical simulations of a homogeneous spatial Poisson process are conducted in MATLAB. The simulations consist in generating random point patterns with a prescribed average density, inside a rectangular observation window identical to the experimental field of view. Ten average densities are prescribed, spanning the range of experimental excavate density $c$ measured. 200 realisations are performed for each density. Results are shown in Fig. S10-S11.

A homogeneous spatial Poisson process is characterised by $g(r) = 1$, $S(k) = 1$ and a number variance $\sigma_N$ scaling as $R^2$. The distribution of its Voronoi areas $A$ follows a gamma distribution, while the distribution of its nearest neighbour distance $d_{nn}$ follows a Weibull distribution with a shape factor $k$ of 2 [5, 6]. The probability density function of a gamma distribution with shape factor $\alpha$ and scale factor $\theta$ is:

$$f_\gamma(x) = \frac{1}{\Gamma(\alpha)\theta^\alpha} x^{\alpha-1} e^{-x/\theta} \tag{S6}$$

for $x \geq 0$, with $\Gamma(\alpha)$ the gamma function. The probability density function of a Weibull distribution with shape factor $k$ and scale factor $\lambda$ is:

$$f_{\text{Weibull}}(x) = \frac{k}{\lambda} \left(\frac{x}{\lambda}\right)^{k-1} e^{(-x/\lambda)^k} \tag{S7}$$

for $x \geq 0$.

#### C. Predator-prey model

To model the predator-prey dynamics, we use a modified version of Lotka-Volterra equations. We introduce a population of resistant bacteria not consumed by excavates ($b_r$) and a population of nutrients ($n^u$), alongside the preey (edible bacteria consumed by excavates $b_e$) and the predators (excavates $c$). The assumptions are that nutrients $n^u$ are consumed by both $b_e$ and $b_r$ at rates proportional to $b_e \cdot n^u$ and $b_r \cdot n^u$. The edible bacteria population $b_e$ grows by consuming nutrients at a rate $\beta_e \cdot b_e \cdot n^u$ and is consumed by excavates at a rate $\alpha \cdot k \cdot b_e \cdot c$. The excavates resistant bacteria population $b_r$ grows by consuming nutrients at a rate $\beta_r \cdot b_r \cdot n^u$ but is not consumed by excavates. The excavates population $c$ grows by consuming edible bacteria $b_e$ at a rate $k \cdot b_e \cdot c$ and dies of starvation at a rate $\delta \cdot c$. Each sub-population of excavates (feeding, swimming, and foraging where relevant) is assumed to behave similarly. In the consumption of edible bacteria $b_e$ by excavates, $\alpha$ accounts for the number of bacteria needed for an excavate to divide. For simplicity, nutrients are counted in the same units as bacteria. The dynamics of the populations are thus described by the following set of differential equations:

$$\frac{\partial n^u}{\partial t} = -(\beta_e \cdot b_e + \beta_r \cdot b_r) \cdot n^u \tag{S8}$$

$$\frac{\partial b_e}{\partial t} = \beta_e \cdot b_e \cdot n^u - \alpha \cdot k \cdot b_e \cdot c \tag{S9}$$

$$\frac{\partial b_r}{\partial t} = \beta_r \cdot b_r \cdot n^u \tag{S10}$$

$$\frac{\partial c}{\partial t} = k \cdot b_e \cdot c - \delta \cdot c \tag{S11}$$

| | *Jakoba libera* | *Kipferlia bialata* | *Reclinomonas americana* |
|---|---|---|---|
| $\beta_e$ (bact$^{-1}$ day$^{-1}$) | $3.5 \times 10^{-5}$ | $2 \times 10^{-5}$ | $35 \times 10^{-5}$ |
| $\beta_r$ (bact$^{-1}$ day$^{-1}$) | $1 \times 10^{-5}$ | $0.5 \times 10^{-5}$ | $10 \times 10^{-5}$ |
| $k$ (bact$^{-1}$ day$^{-1}$) | $0.6 \times 10^{-5}$ | $8 \times 10^{-5}$ | $4 \times 10^{-5}$ |
| $\alpha$ (bact cell$^{-1}$) | 200 | 75 | 60 |
| $\delta$ (day$^{-1}$) | 0.09 | 0.02 | 0.05 |
| $n^u(0)$ (bact) | $1.7 \times 10^5$ | $4 \times 10^4$ | $0.55 \times 10^5$ |
| $b_e(0)$ (bact) | 1.5 | 40 | 100 |
| $b_r(0)$ (bact) | $2 \times 10^3$ | $7 \times 10^3$ | $7 \times 10^3$ |
| $c(0)$ (cells) | 23 | 2 | 1.5 |

Table S1. Parameters used to fit the predator-prey model to the growth curves of the three typical excavates in Fig. 4a-c.

### II Ultrastructure Expansion Microscopy (U-ExM)

Morphology measurements of *J. libera*, *K. bialata*, and *R. americana* were made manually from the 3D reconstruction of the U-ExM images (see Methods) in DragonFly 3D World, see Table S2. The body length corresponds to the distance between the anterior and the posterior end of the cell (along the major axis of the groove), the width to the direction along the minor axis of the groove, and the height to the direction perpendicular to the groove. These measurements are consistent with the literature [7, 8].

Multiple spots of DNA consistently appear at the anterior end of all imaged *J. libera* cells ($n = 32$ cells), on top of the spots visible at the posterior end, see Fig. S1, potentially indicating the presence of ingested bacteria or endosymbionts. DNA spots were not observed at the anterior end of the cells for *K. bialata* or *R. americana*.

| | *Jakoba libera* ($n = 12$ cells) | *Kipferlia bialata* ($n = 5$ cells) | *Reclinomonas americana* ($n = 8$ cells) |
|---|---|---|---|
| Body length (µm) | $8.0 \pm 1.3$ | $9.8 \pm 1.7$ | $7.1 \pm 1.6$ |
| Body width (µm) | $4.4 \pm 0.3$ | $6.2 \pm 1.0$ | $4.5 \pm 1.3$ |
| Body height (µm) | $2.8 \pm 0.9$ | $4.7 \pm 0.8$ | $2.8 \pm 0.8$ |
| Anterior flagellum length (µm) | $13.4 \pm 1.3$ | $11.7 \pm 1.6$ | $11.1 \pm 3.0$ |
| Posterior flagellum length (µm) | $12.3 \pm 2.3$ | $14.9 \pm 1.4$ | $12.2 \pm 1.8$ |

Table S2. Morphology measurements of typical excavates from Ultrastructure Expansion Microscopy (U-ExM) images (see Methods). Values have been corrected with the expansion factor, and correspond to the average ± standard deviation for the $n$ cells of each species.

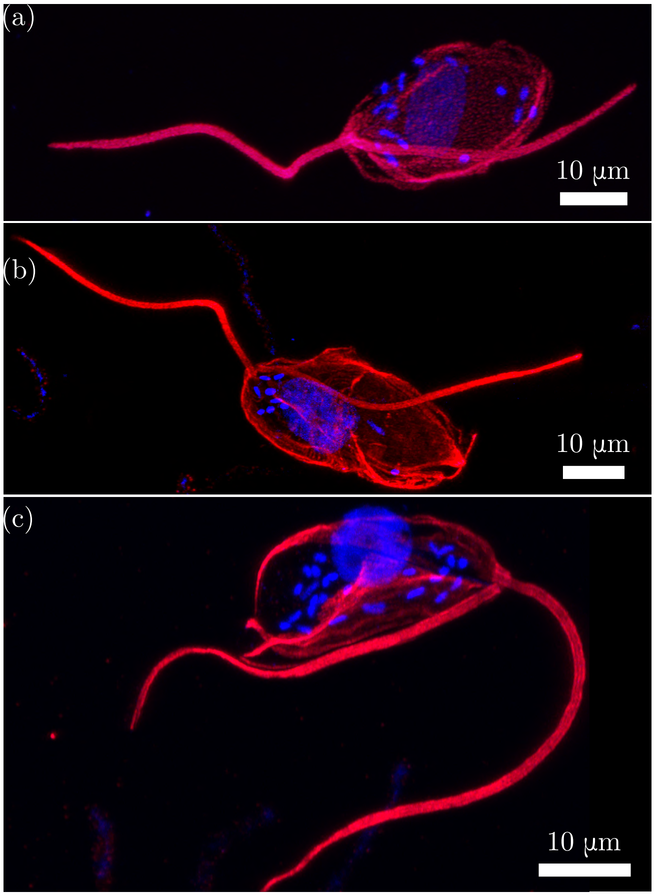


Figure S1. **U-ExM images of *Jakoba libera*.** (a-c) Maximum intensity projection of *J. libera* cells expanded by Ultrastructure Expansion Microscopy (U-ExM), see Methods. Cells were expanded (a) 4.5×, and (b-c) 4.0×. Tubulin (red) and DNA (blue) are visible. An accumulation of bacteria is visible at the anterior end of each cell.

### III Bacteria morphology

Bacterial species composition varies significantly among the three typical excavates cultures. The samples were originally collected from different locations and different natural environments (*J. libera* is marine aerobic, *K. bialata* is marine anaerobic, and *R. americana* is freshwater aerobic). Snapshots of the bacteria and their distributions close to the bottom surface are shown in Fig. S2.

The bacteria in *J. libera* cultures are long, filamentous, and mostly sessile, and form biofilms. The bacteria in *K. bialata* cultures are highly diverse in shape, size and motility. They are very numerous both near surfaces and throughout the entire water column. Due to this abundance and shape diversity, the true density of bacteria in *K. bialata* cultures is difficult to resolve accurately. The bacteria in *R. americana* cultures are typically 1 to 2 µm long, rod-shaped, and sessile.

### IV High-speed characterisation of the *J. libera* swimming behaviour

High-speed imaging of swimming *J. libera* cells was conducted in the fluid bulk far from surfaces (see Methods). An example trajectory showing straight swimming in one direction, followed by a turn, and then swimming in a new direction is shown in Fig. 2h (also Movie S3). This excavate was recorded at 500 fps with 101× magnification, and remained in the field of view for $\approx 4$ s. This high magnification and frame rate allow for high-resolution tracking of its motion, see Fig. S3a.

This type of trajectory reveals further features of the dynamics not detectable in lower-frame-rate videos (such as those in Fig. 3a). Here the swimming speed of the 'run' is around $14\,\mu\mathrm{m\,s^{-1}}$, see black line in Fig. S3c, before a turn happens at $t \approx 2.65$ s. These 'runs' are not perfectly straight, but display large oscillations orthogonal to the swimming direction (Fig. S3a). These oscillations are due to the self-rotation of the excavate around its long-axis, which produces the helical trajec-

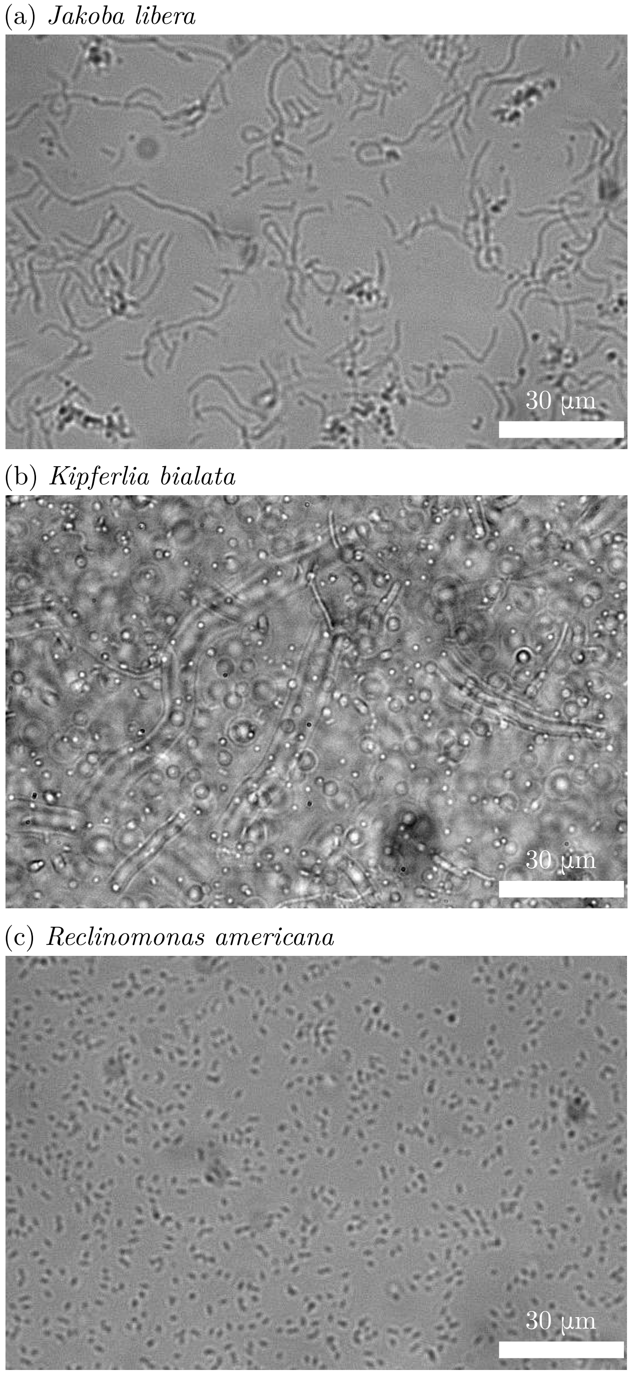


Figure S2. **Endemic bacteria in cultures of typical excavates.** Images of the endemic bacteria in cultures of (a) *Jakoba libera*, (b) *Kipferlia bialata*, and (c) *Reclinomonas americana*. Images were taken on the bottom surface with a 40× objective. Measured bacterial densities $b$ are respectively (a) $b = 6.6 \times 10^4$ bact mm$^{-2}$, (b) $b = 4.7 \times 10^4$ bact mm$^{-2}$, and (c) $b = 6.2 \times 10^4$ bact mm$^{-2}$. Age of the cultures is respectively (a) $t = 3$ days, (b) $t = 8$ days, and (c) $t = 6$ days.

tories. This is associated with a rolling frequency of approx $f_{\mathrm{roll}} = 1/\delta t \approx 1.5$ Hz, along with an amplitude of $\approx 1.5$ µm. Zooming in even further, we detect another oscillation frequency (inset in Fig. S3c) which corresponds to the back-and-forth motion caused by the beating of the anterior flagellum. Its beating frequency $f$ can thus be measured as $f = 1/\Delta t \approx 15$ Hz, along with an amplitude of $\approx 0.5$ µm, consistent with previous measurements made by Suzuki-Tellier *et al.* [7].

### V High-speed characterisation of the *Jakoba libera* foraging behaviour

A higher sampling rate (100 fps) dataset of foraging *J. libera* cells allows us to resolve the dynamics at short times, which revealed a second diffusive regime at short times ($t < 0.2\,s$), see Fig. S4. This regime is characterised by a higher diffusivity than the diffusive regime at long times, at least when prey availability is high,

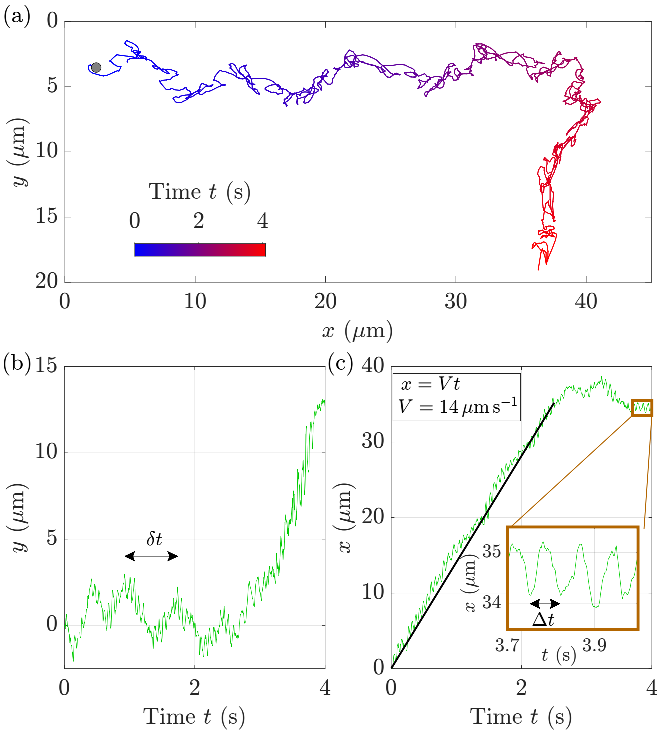


Figure S3. **Zoom in on the swimming behaviour of excavate *Jakoba libera*.** (a) Typical swimming trajectory of excavate *J. libera* in the bulk. Same trajectory as in Fig. 2h and Movie S3, but rotated by 10°. (b-c) Temporal evolution of the positions $y$ (b) and $x$ (c) of the trajectory shown in (a). From the first part of the trajectory before the turn ($t < 2.65$ s), the swimming velocity can be measured with a linear fit $x = Vt$. A velocity of $V = 14$ µm s$^{-1}$ is found. From the small oscillations in $x$ or $y$, see inset in (c), the beating frequency of the anterior flagellum can be extracted: $f = 1/\Delta t \approx 15$ Hz. From the large oscillations in $y$, the 'rolling' frequency of the excavate can be measured: $f_{\mathrm{roll}} = 1/\delta t \approx 1.5$ Hz. This rolling time $\delta t$ corresponds to the time needed for the excavate to do one full rotation on itself, i.e., to travel along one screw pitch of its helical trajectory.

with $D \approx 5.9\,\mu m^2\,s^{-1}$. A transition appears between $t = 0.2 - 3\,s$, linking the two diffusive regimes. At long times ($t > 3\,s$), the diffusive behaviour observed in Fig. 2b (blue diamonds) is recovered, characterised by a diffusion coefficient of $D \approx 0.75\,\mu m^2\,s^{-1}$.

These two distinct diffusive regimes can be interpreted thus: the excavates diffuse around a singular bacterial hotspot at short times, and diffuse at longer times by hopping between different hotspots. The critical transition timescale, here $t = 0.2 - 3\,s$, would then correspond to the residence time of the cells around one hotspot.

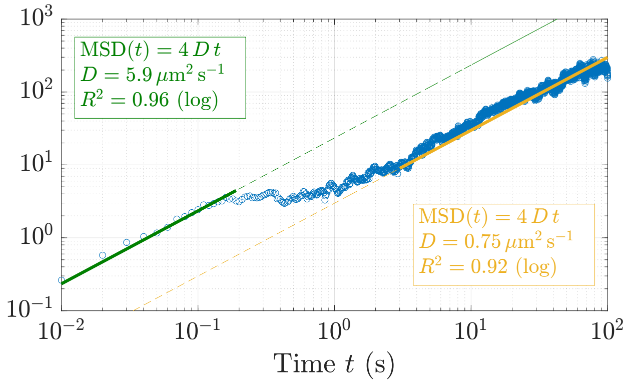


Figure S4. **High-speed characterisation of the foraging behaviour of excavate *Jakoba libera*.** Mean squared displacement (MSD) as a function of time $t$ for a 5-day old population of *J. libera* recorded near the bottom surface, at 100 fps with a 40× objective.

### VI Motility of *Kipferlia bialata*

*K. bialata* exhibits a similar motility to *J. libera*. *K. bialata* cells perform feeding, swimming, and foraging (see Fig. S5). Their swimming velocity ranges from 5 to $15\,\mu m\,s^{-1}$, similar to *J. libera* but much lower than *R. americana*.

Changes in the excavates motility are noticeable over the culture cycle, with an increase in both the swimming population and the diffusion coefficient of the foraging sub-population when food availability decreases. The swimming population $c_{swim}/c$ increases from 0 % to $\approx$ 2 %, while the diffusion coefficient $D$ rises from $\approx 1\,\mu m^2\,s^{-1}$ to 10-20 $\mu m^2\,s^{-1}$. The mean velocity $V$ of the swimming population remains constant around $11\,\mu m\,s^{-1}$. These trends match *J. libera* behavioural changes, although our data on *K. bialata* is not enough to conclude with certainty (short trajectories of only 1 biological replicate, due to difficulties in culturing this species).

### VII Characterisation of *Jakoba libera* trajectories

Single-cell swimming trajectories were used to resolve and classify *J. libera* motility into distinct modes of swimming, foraging or feeding. Initially, each unique trajectory was categorised as one of the three motility modes (swimming, foraging and feeding). The distance travelled $d_{travel}$, i.e., the distance between the start and end points of the trajectory, was a good indicator of whether a cell is feeding or not, as cells are attached to the surface while feeding. An excavate is classed as 'feeding' if $d_{travel} \leq 8\,\mu m$, based on the distributions of distance travelled. With a median velocity $v_{med}$ higher than $6\,\mu m\,s^{-1}$, an excavate is considered 'swimming'. To avoid false positives, strict criteria are used for the 'foraging' population: an excavate needs to satisfy both $d_{travel} > 8\,\mu m$ and $\max(v) < 6\,\mu m\,s^{-1}$ to be considered 'foraging'. See Fig. S6 for the application of these criteria.

To further resolve the transitions between different motility modes (such as occurring within a single trajectory), we performed an additional analysis step. This step was also necessary to precisely measure the fraction of swimming excavates. Thresholds based on instantaneous velocity $v$ (see Methods) gave the best result. We use the velocity $v$ with a threshold of $6\,\mu m\,s^{-1}$ to separate swimming tracks from foraging and feeding ones, see Fig. S7a for a thresholded view of the typical tracks shown in Fig. 3a. The temporal evolution of $v$ for four different trajectories is also shown in Fig. S7b with the threshold plotted, including two trajectories showing transitions from foraging to swimming behaviour (green and orange), one showing only swimming (yellow) and one only foraging behaviour (blue).

### VIII Excavates competing for a feeding spot

Interactions between excavates were also observed, and hint at competition for limited resources. In one example, *J. libera* cells were navigating around the same feeding spot on the bottom surface, see Fig. S8. Initially, excavate A is attached to a bacterial biofilm in a feeding state (teal), while excavate B is foraging along the surface (brown). Later, B encounters A and takes its place, see Fig. S8c. They immediately swap behaviours, with B switching to feeding mode while A switches to foraging. Surprisingly, the now-foraging excavate A then circles back to its original feeding spot after a couple of minutes, and displaces B, whereupon they again immediately swap behaviours. Eventually, B leaves the spot and diffuses along the surface, presumably to look for a new feeding location.

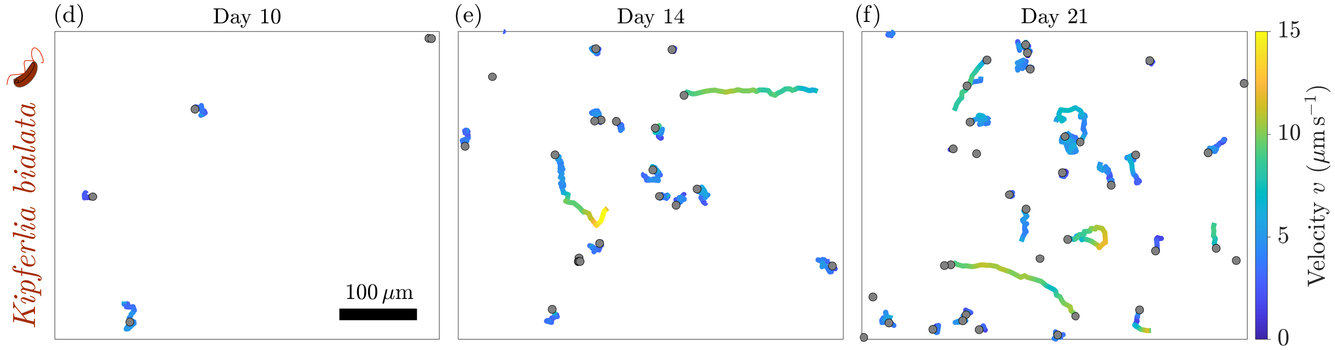


Figure S5. **Motility of *Kipferlia bialata* over time.** (a-c) Trajectories of *K. bialata* imaged over 20 s near the bottom surface, in (a) early growth-phase ($t = 10$ days), (b) late growth-phase ($t = 14$ days), or (c) saturation phase ($t = 21$ days). Grey points mark the starting point of the tracks. Note that the trajectories shown here are shorter (20 s) than the ones displayed in Fig. 4d-i for *Jakoba libera* (10 min) and *Reclinomonas americana* (1.5 min).

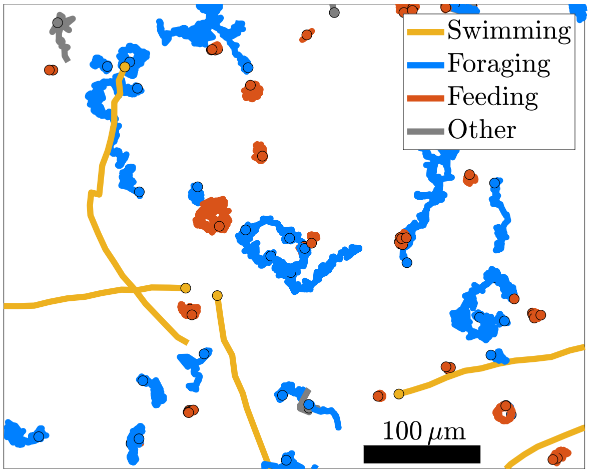


Figure S6. **Trajectory classification for *Jakoba libera*.** Trajectories of *J. libera* close to the bottom surfacein an intermediate dense bacteria suspension ($t = 7$ days). Each trajectory is coloured according to the (predominant) behavioural state, yellow for swimming ($v_{\mathrm{med}} > 6\,\mu\mathrm{m\,s^{-1}}$), blue for foraging ($d_{\mathrm{travel}} > 8\,\mu\mathrm{m}$ and $\max(v) < 6\,\mu\mathrm{m\,s^{-1}}$), orange for feeding ($d_{\mathrm{travel}} \leq 8\,\mu\mathrm{m}$), and grey for the trajectories not fulfilling any criteria. Circled points mark the starting point of the tracks.

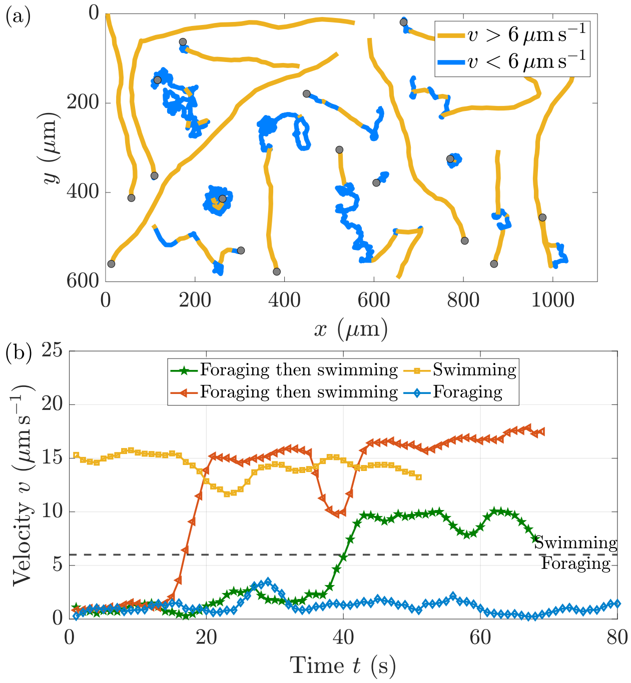


Figure S7. **Identification of motility transitions in excavate *Jakoba libera* trajectories.** (a) Typical trajectories near surfaces, coloured with their instantaneous velocity $v$ using a threshold of $6\,\mu\mathrm{m\,s^{-1}}$. Thresholded version of Fig. 3a. Grey points mark the starting point of the tracks. (b) Temporal evolution of $v$ for 4 trajectories: 2 exhibiting a foraging behaviour before transitioning to swimming (green and red), 1 swimming trajectory (yellow), and 1 foraging trajectory (blue).

### IX Feeding current of a loricate *Reclinomonas americana* cell

To estimate the range of the feeding current generated by a loricate *R. americana*, sessile bacteria naturally present in the suspension were manually tracked as a proxy for the flow field (see Movie S7). Trajectories show that bacteria close enough to the bottom-right side of the cell are attracted by the feeding current and repelled when on the top-left side, see Fig. S9a. Their instantaneous velocities $v_{\mathrm{flow}}$ (smoothed with a moving average on a 1.5 s-window) locally reach a maximum around $15\,\mu\mathrm{m\,s^{-1}}$ while passing next to the excavate. $v_{\mathrm{flow}}$ decreases as the distance $r$ to the excavate increases, see Fig. S9b. No significant velocity is measured beyond $r \approx 75\,\mu\mathrm{m}$. Note that the feeding current seems to be restricted in angle, and looks very similar to a Stokeslet [9, 10]. A power-law fit $v_{\mathrm{flow}} = A_0 r^n$ of our data results produces a $n = -0.8$ exponent (see orange

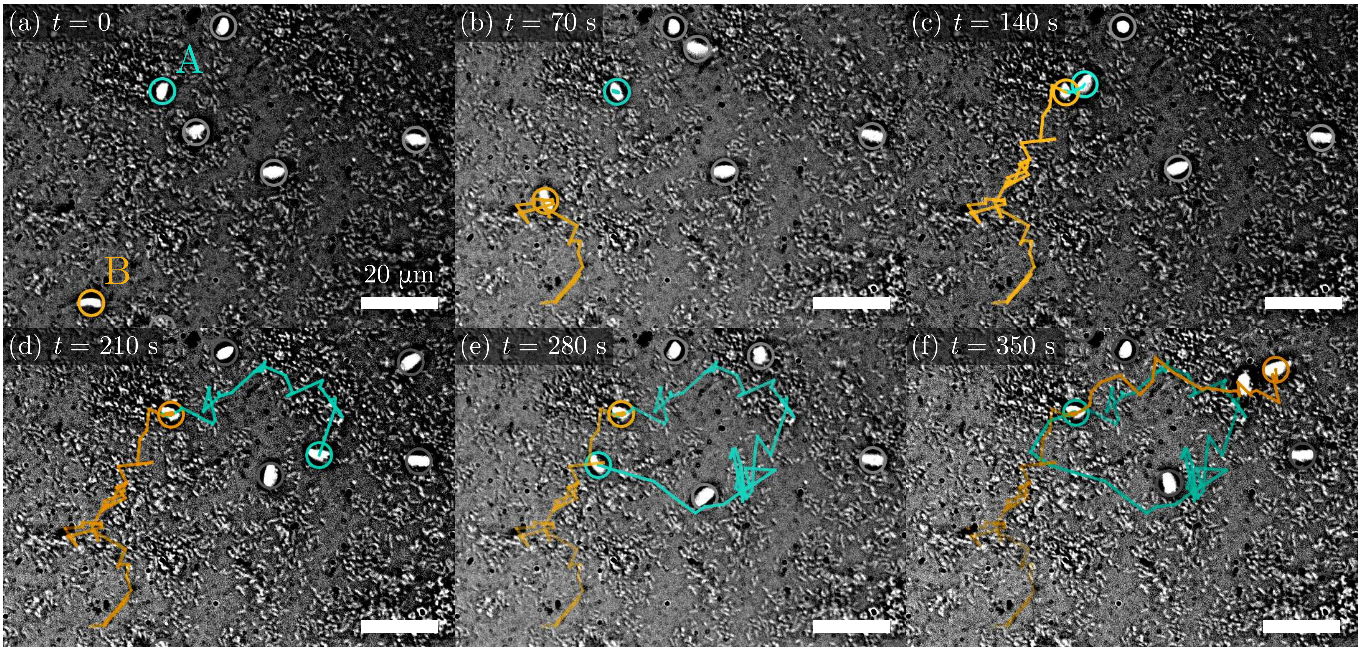


Figure S8. **Two excavates *Jakoba libera* competing for a feeding spot.** Timelapse of excavates *J. libera* feeding and foraging on bacterial biofilms, see Movie S6. The trajectories of two excavates are highlighted in teal (A) and brown (B). These two excavates switch between foraging and feeding behaviour when they swap places at a specific interesting feeding spot. Movie recorded on the bottom surface of a 6-day old culture, at a frame rate of 0.35 fps with a 40× objective. Scalebars correspond to 20 µm.

line in Fig. S9b), which is compatible with the $1/r$ decay expected for a Stokeslet. For reference, an exponential function $v_{\text{flow}} = v_0 e^{-r/r_0}$ has also been fitted to the data (black line).

### X Variations of spatial statistics with excavates density

The distribution of excavates on the surface can be characterised by the Voronoi polygon area $A$, the nearest neighbour distance $d_{nn}$, the pair distribution function $g(r)$, the structure factor $S(k)$, and the number variance $\sigma_N$ (see Methods). Evolution of these spatial statistics indicators is now explored as a function of the density of bacteria $b$, the density of excavates $c$ and the prey availability $\gamma = b/c$. The scaling with respect to $c$ was found to be the most informative, and is shown in Fig. S10-S11 for *J. libera* (left), *R. americana* (right), and the Poisson simulations (right, see Methods). The Poisson simulations provides the baseline for a random distribution of points in a finite observation window, and helps us in identifying trends that are characteristic of the excavates populations.

For *J. libera*, the short-range attraction seen in the agglomerated data (see Fig. 6) is particularly important at low excavate density $c$. It is characterised by a huge peak in $g(r)$ as well as higher probabilities for small values of $d_{nn}/\bar{d}$ and $A/\bar{d}^2$. At high $c$, the long tails of the two distributions are clear deviations from the distributions expected by a Poisson process. For *R. americana*, no significant evolution with $c$ is observed on the curves of $P(A/\bar{d}^2)$, $P(d_{nn}/\bar{d})$ or $g(r)$.

For both species, the average structure factor differs from $S(k) = 1$, the expected value of a random spatial distribution (as confirmed by the Poisson simulations). At very low wavenumber ($k < 1 \times 10^{-2}\,\mu\text{m}^{-1}$), an increase of $S(k)$ is observed for both species. However, at intermediate wavenumber ($1 \times 10^{-2}\,\mu\text{m}^{-1} > k > 5 \times 10^{-2}\,\mu\text{m}^{-1}$), the structure factor of *R. americana* is lower than the baseline, unlike that of *J. libera*. Changes in the excavate density do not seem to influence $S(k)$, but it is difficult to conclude with certainty due to the high noise in the data.

The variance in numbers $\sigma_N$ is clearly affected by the finite-size of the field of view, since the Poisson simulations deviate from the baseline at high values of $R$ ($R > 2 \times 10^2\,\mu\text{m}$). The experimental data of the two excavates show the same sharp decrease at high $R$. Nevertheless, these curves are very informative at lower value of $R$. Firstly, the excavate density $c$ has a big impact on the dynamics of $\sigma_N$ for both species. For *J. libera*, $\sigma_N/R^2$ increases as $R$ increases for high $c$, but decreases for low $c$ (except for the lowest bin). For *R. americana*, $\sigma_N/R^2$ decreases as $R$ increases for high and low $c$, but increases for intermediate densities. These decreases with $R$, observed for both species but especially for *R. americana*, is indicative of weak hyperuniformity [11, 12].

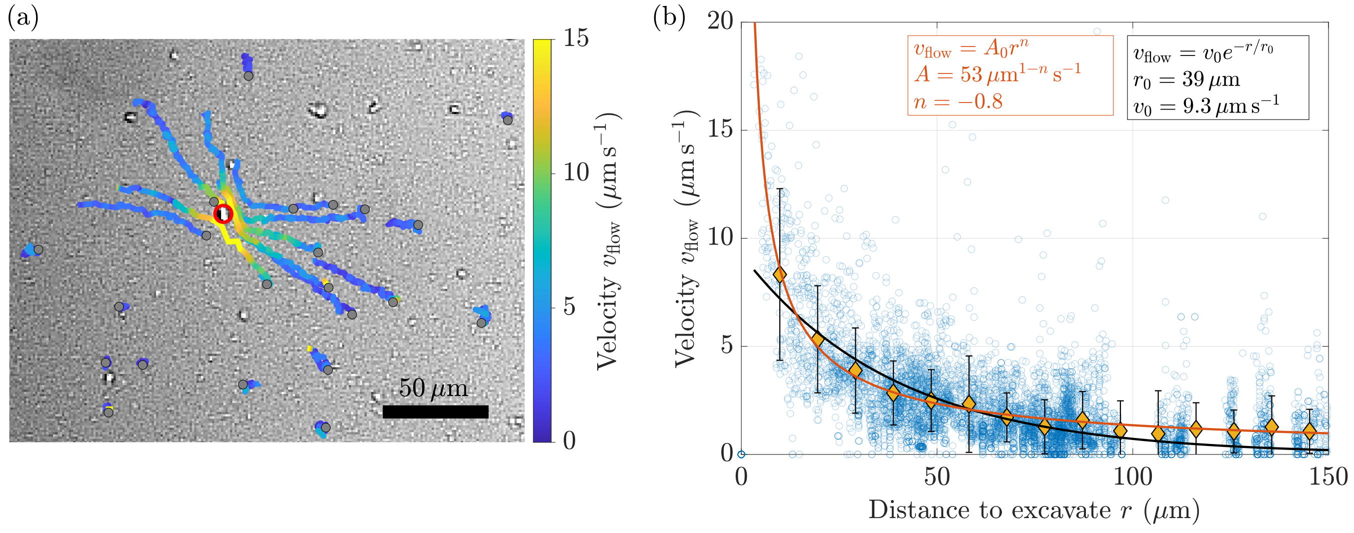


Figure S9. **Bacteria displaced by the feeding current of a loricate *Reclinomonas americana* cell.** (a) Trajectories of sessile bacteria around a feeding *R. americana* as proxy for the feeding flow field (34-day old culture), coloured by their instantaneous velocity $v_{\text{flow}}$ (see Movie S7). Grey points mark the starting point of the tracks. Red circle: location of the excavate. The movie is recorded with a 10× objective at 6.8 fps. (b) Evolution of $v_{\text{flow}}$ as a function of the distance $r$ ($n = 2$ cells). Experimental data (blue circles) are shown along their binning in $r$ (yellow diamonds). Lines: Fit with a power-law (orange) or an exponential function (black).


[1] J. Schindelin, I. Arganda-Carreras, E. Frise, V. Kaynig, M. Longair, T. Pietzsch, S. Preibisch, C. Rueden, S. Saalfeld, B. Schmid, *et al.*, "Fiji: an open-source platform for biological-image analysis," *Nature Methods*, vol. 9, no. 7, pp. 676–682, 2012.

[2] J.-Y. Tinevez, N. Perry, J. Schindelin, G. M. Hoopes, G. D. Reynolds, E. Laplantine, S. Y. Bednarek, S. L. Shorte, and K. W. Eliceiri, "Trackmate: An open and extensible platform for single-particle tracking," *Methods*, vol. 115, pp. 80–90, 2017.

[3] G. Voronoi, "Nouvelles applications des paramètres continus à la théorie des formes quadratiques. deuxième mémoire. recherches sur les parallélloèdres primitifs.," *Journal für die reine und angewandte Mathematik (Crelles Journal)*, vol. 1908, no. 134, pp. 198–287, 1908.

[4] T. C. Day, S. S. Höhn, S. A. Zamani-Dahaj, D. Yanni, A. Burnetti, J. Pentz, A. R. Honerkamp-Smith, H. Wioland, H. R. Sleath, W. C. Ratcliff, *et al.*, "Cellular organization in lab-evolved and extant multicellular species obeys a maximum entropy law," *Elife*, vol. 11, p. e72707, 2022.

[5] P. J. Diggle, *Statistical analysis of spatial and spatio-temporal point patterns*. CRC press, 2013.

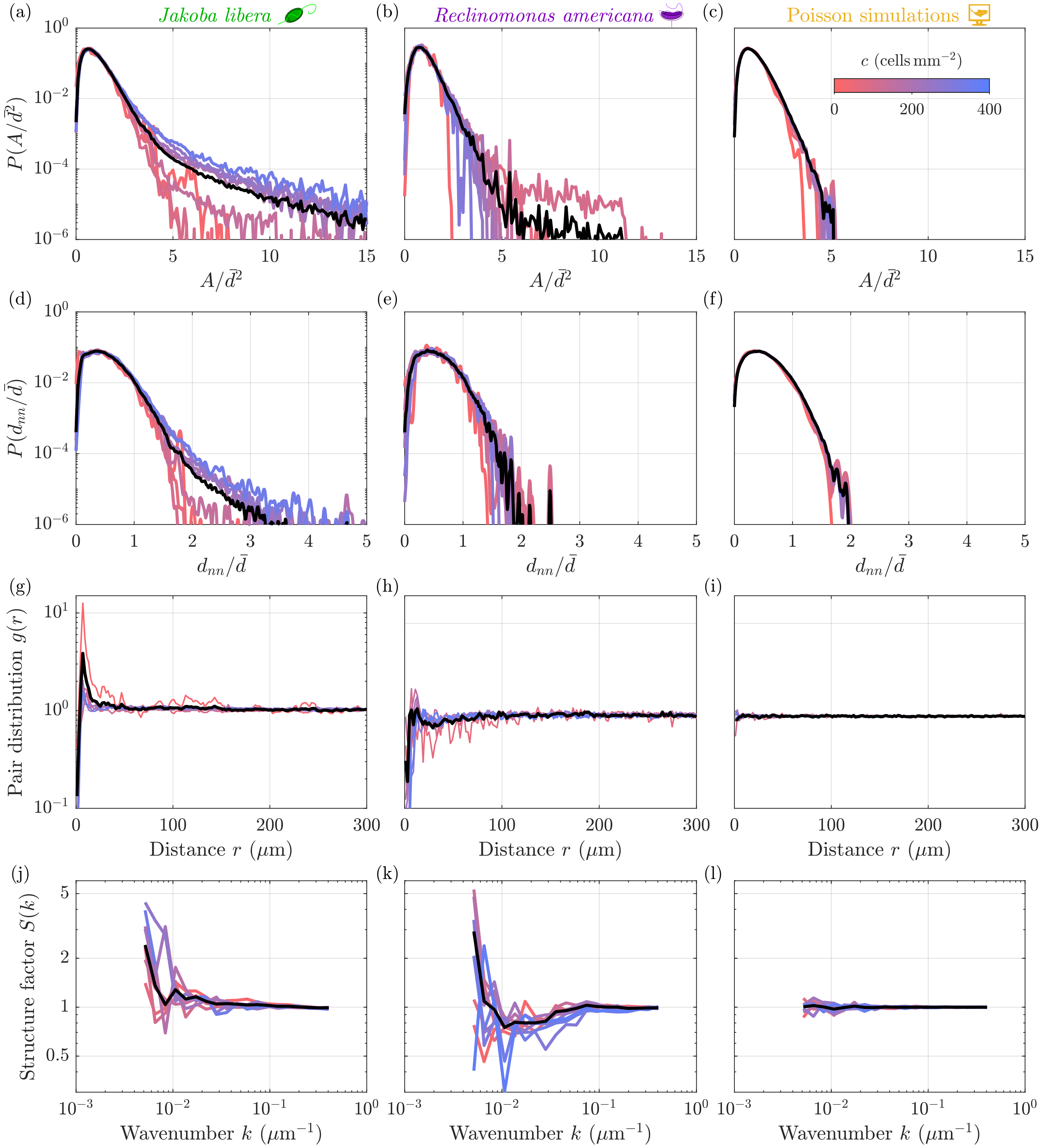


Figure S10. **Spatial statistics dependence on the excavates density** $c$**.** Variations of (a-c) the probability distributions of normalised Voronoi area $P(A/\bar{d}^2)$, (d-f) the probability distributions of normalised distance to nearest neighbour $P(d_{nn}/\bar{d})$, (g-i) pair distribution functions $g(r)$, and (j-l) structure factors $S(k)$ with the excavates density $c$ (see Methods). Data shown for (a,d,g,j) *Jakoba libera*, (b,e,h,k) *Reclinomonas americana*, and (c,f,i,l) numerical simulations with a homogeneous spatial Poisson process (see Methods). Colourscale is identical for each panel. Black curves represent the ensemble average.

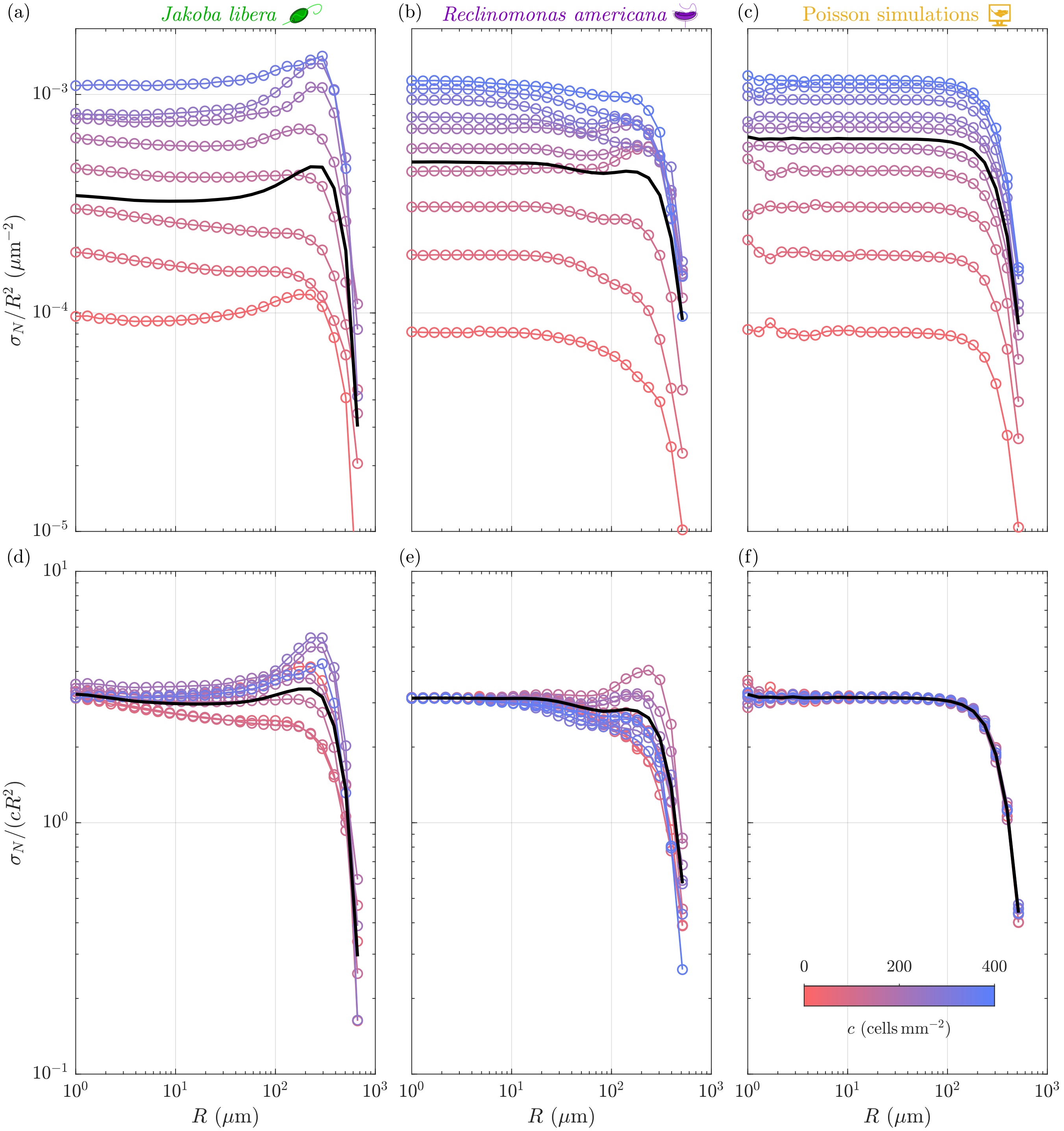


Figure S11. **Variations of number variance $\sigma_N$ as a function of the window size $R$, with excavates density $c$.** Variations of (a-c) the number variance normalised by $R^2$ – $\sigma_N/R^2$ – and (d-f) the number variance further normalised by $c$ – $\sigma_N(R)/(cR^2)$ – with the excavates density $c$ (see Methods). Data shown for (a,d) *Jakoba libera*, (b,e) *Reclinomonas americana*, and (c,f) numerical simulations with a homogeneous spatial Poisson process (see Methods). Colourscale is identical for each panel. Black curves represent the ensemble average.

### XI Legends for Supplementary Movies

Movie S1 – High-speed movie of a feeding *Jakoba libera* cell attached to a surface.

Movie S2 – High-speed movie of two foraging *Jakoba libera* cells along surfaces.

Movie S3 – High-speed movie of a swimming *Jakoba libera* cell in the bulk.

Movie S4 – High-speed movie of a feeding, loricate *Reclinomonas americana* cell attached to a surface.

Movie S5 – High-speed movie of a swimming *Reclinomonas americana* cell in the bulk.

Movie S6 – Competition between two *Jakoba libera* for a feeding spot in a bacterial biofilm.

Movie S7 – Feeding current of a loricate *Reclinomonas americana* cell, visualised via the displacement of sessile bacteria suspended around the cell.
